# CHOIR improves significance-based detection of cell types and states from single-cell data

**DOI:** 10.1101/2024.01.18.576317

**Authors:** Cathrine Sant, Lennart Mucke, M. Ryan Corces

**Affiliations:** Gladstone Institute of Neurological Disease, Gladstone Institutes, San Francisco, CA, USA; Neuroscience Graduate Program, University of California, San Francisco, San Francisco, CA 94158, USA; Department of Neurology and Weill Institute for Neurosciences, University of California, San Francisco, San Francisco, CA 94158, USA

## Abstract

Clustering is a critical step in the analysis of single-cell data, as it enables the discovery and characterization of putative cell types and states. However, most popular clustering tools do not subject clustering results to statistical inference testing, leading to risks of overclustering or underclustering data and often resulting in ineffective identification of cell types with widely differing prevalence. To address these challenges, we present CHOIR (<u>c</u>lustering <u>h</u>ierarchy <u>o</u>ptimization by iterative random forests), which applies a framework of random forest classifiers and permutation tests across a hierarchical clustering tree to statistically determine which clusters represent distinct populations. We demonstrate the enhanced performance of CHOIR through extensive benchmarking against 14 existing clustering methods across 100 simulated and 4 real single-cell RNA-seq, ATAC-seq, spatial transcriptomic, and multi-omic datasets. CHOIR can be applied to any single-cell data type and provides a flexible, scalable, and robust solution to the important challenge of identifying biologically relevant cell groupings within heterogeneous single-cell data.

## INTRODUCTION

Cellular heterogeneity is a fundamental characteristic of complex biological systems, which are functionally dependent on the concerted action of many distinct, specialized cell populations. Single-cell technologies make this cellular heterogeneity accessible, enabling the recognition and characterization of distinct cell populations. Therefore, partitioning cells into empirical groups (or clusters) representing biologically distinguishable cell types or cell states is an important objective in the analysis of single-cell data. Whether the clusters obtained from such analyses are reliable proxies for biologically distinct cell populations materially affects the ability of subsequent analysis steps, such as differential expression analyses, to yield meaningful biological insights. Underclustering, in which multiple distinct populations are grouped into the same cluster, may obscure subtle but potentially impactful differences between cell subtypes or cell states, whereas overclustering, in which cluster divisions are driven by noise rather than true biological signal, may lead to the misguided investigation of spurious new cell types or states.

A variety of clustering approaches have been developed for use on single-cell data. However, in many biological systems studied with single-cell approaches, the prevalence of specific putative cell types can differ by orders of magnitude^1^. This wide range presents a challenge for many clustering methods, including Louvain^2^ and Leiden^3^ clustering, which often perform best when cluster sizes are relatively similar^4^. Although some tools have been specifically developed to identify rare cell types^5,6^, the simultaneous and reliable identification of both rare and abundant cell populations remains a major challenge.

In practice, clustering of single-cell data has become a very manual process that is impacted by subjective decisions based, in part, on biological intuition, which varies from user to user. More specifically, the granularity of clustering results is often greatly influenced by parameter values selected by the user, such as the “resolution” parameter in Louvain clustering, which greatly influences the number of clusters and enables the algorithm to find smaller clusters of tightly connected nodes at higher resolution values. Users often manually assess multiple clustering results based on toggling of these parameters before selecting one that matches their intuition. Moreover, it has become commonplace to use this initial clustering result as a starting point for manual curation by merging or dividing (“subclustering”) certain selected clusters. However, gauging the extent to which clusters should be merged or subdivided is difficult, as many clustering methods will partition data even if they are derived from a single cell population and no biologically meaningful groups are present^4^. Because of the subjective nature of this approach, there are limited safeguards against overclustering or underclustering of the data.

Because the manual assessment of clustering results is neither scalable nor reproducible, methods have been developed to circumvent this process. Methods to estimate the number of “true” clusters in the data, as well as cluster stability methods^7,8^, can help assess the reproducibility of clusters across subsamples or parameters, but often overestimate the number of clusters present^9^ and do not account for statistical uncertainty. Similarly, reference-based clustering methods have been developed to automate the annotation of new datasets, leveraging the cell type markers identified in previous datasets. However, these supervised approaches are limited to—and inherently biased by—the annotations used as input, which are largely based on the same flawed principle of leveraging biological intuition to determine proper clustering. Because of these limitations, such methods may fail to identify novel cell types or states.

Another potential limitation of existing single-cell clustering methods is that most have been optimized specifically for single-cell RNA sequencing (scRNA-seq) data. However, the rise of additional single-cell data types, including from multi-omic approaches, enables concerted insights into cellular phenotypes across multiple modalities, such as mRNA, chromatin accessibility, and surface protein abundance. To fully leverage this expanding landscape of technologies, computational tools must be adaptable across multiple data types and be able to perform joint multi-modal analyses.

Thus, there is a need for single-cell clustering methods that can reliably detect rare as well as common cell types across single– and multi-omic platforms, while neither overclustering nor underclustering the data. In addition, the ideal clustering method would reduce the chance of spurious discoveries by incorporating statistical inference and minimizing the reliance on user-supplied parameters with substantial effects on clustering output. Imposing such algorithmic guardrails could improve one’s confidence in the biological relevance of single-cell results and save time and effort that would otherwise be spent applying intuition to manually curate clustering results. To address these needs, we have developed CHOIR (<u>c</u>luster <u>h</u>ierarchy <u>o</u>ptimization by iterative random forests), a user-friendly R package for clustering single-cell data of any modality. CHOIR integrates seamlessly with popular single-cell sequencing software packages such as Seurat^10^, SingleCellExperiment^11^, ArchR^12^, and Signac^13^. It uses a hierarchical permutation test approach based on random forest classifier predictions to identify clusters representing distinct cell types or states with increased levels of accuracy and ease, compared to a broad range of available clustering algorithms and to ground truths established or estimated by other methodologies.

## RESULTS

### Overview of the CHOIR algorithm

CHOIR is based on the premise that, if clusters contain biologically different cell types or states, a classifier that considers features present in cells from each cluster should be able to distinguish the clusters with a higher level of accuracy than classifiers trained on randomly permuted cluster labels (**Fig. 1a**). The use of a permutation testing approach allows CHOIR to introduce statistical significance thresholds into the clustering process. Operating as a self-contained clustering algorithm, CHOIR iteratively applies this permutation test approach across the nodes of a hierarchical clustering tree to determine the appropriate extent of each branch of the clustering tree such that the final clusters represent statistically distinct populations across a range of cluster sizes. Random forest classifiers are used because of their flexible applicability to data derived from various distributions. Consequently, CHOIR is applicable to single-cell datasets across diverse technologies, including RNA-seq, ATAC-seq, and proteomics.

**Fig. 1:**
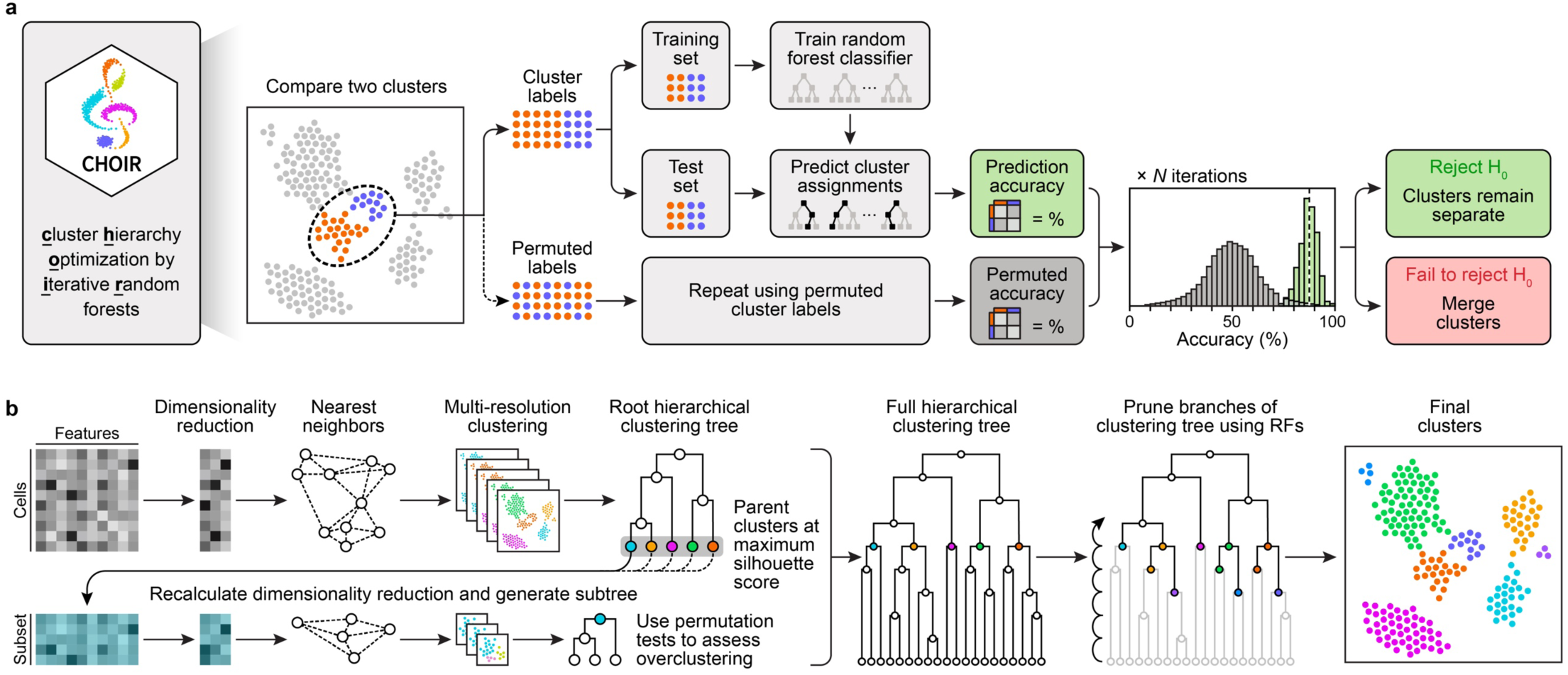
CHOIR is a hierarchical clustering algorithm that uses permutation testing for cluster identification by statistical inference. **a.** Schematic demonstrating how CHOIR identifies clusters that should be merged by applying a permutation test approach to assess the accuracy of random forest classifiers in predicting cluster assignments from a normalized feature matrix. **b.** Schematic demonstrating how CHOIR constructs and iteratively prunes a hierarchical clustering tree using statistical inference to prevent underclustering and overclustering.

CHOIR takes as input a Seurat^10^, SingleCellExperiment^11^, or ArchR^12^ object containing one or more feature matrices (**Supplementary Fig. 1a–b**). Input data should already have undergone quality control and normalization steps. CHOIR proceeds in two main steps (**Fig. 1b**). First, a hierarchical clustering tree is generated spanning the network structure of the data using a “top-down” approach that proceeds from an initial “partition,” in which all cells are in the same cluster, to a partition in which all cells are demonstrably overclustered. To generate this hierarchical clustering tree, the data is iteratively clustered using modularity-based algorithms^2,3^ with increasing resolution parameters. The silhouette score, which measures the similarity of cells to their own clusters compared to other clusters^14^, is assessed at each new level of the emerging tree. When the silhouette score is maximized, the cells from each “parent cluster” at this level are subsetted for re-analysis. For each parent cluster subset, a new set of highly variable features is identified, a new dimensionality reduction is generated, and these cells are then subclustered to generate a hierarchical clustering subtree. For example, when applied to a multi-omic dataset with both RNA-seq and ATAC-seq features from 51,409 human retina cells^15^, CHOIR identifies three parent clusters (**Extended Data Fig. 1a–b, Supplementary Table 1–2**). Each of the resulting subtrees continues to be subdivided until the farthest pair of nearest neighboring clusters is found to be overclustered by the permutation test approach, as explained further below.

Second, this overclustered hierarchical clustering tree is “pruned” using a bottom-up approach to identify a final set of clusters. Starting at the bottom level of the clustering tree, CHOIR identifies sets of clusters that belong to the same composite cluster at the next level of the clustering tree. For each such cluster pair, balanced random forest classifiers are applied to learn the cluster labels and permuted cluster labels. For these permutation test comparisons, the null hypothesis is that the cells belonging to each of the two clusters are no more distinguishable than a chance separation of the cells into two random groups. If this null hypothesis is rejected, the clusters will retain separate identities. If the null hypothesis is not rejected, the clusters will be merged and assume a single identity (**Fig. 1a**). This process is repeated up the branches of the hierarchical clustering tree until a final cluster solution is reached, wherein each cluster can be accurately distinguished from all other clusters (**Fig. 1b**). The combined top-down tree construction and bottom-up branch pruning approach used by CHOIR ensures that the most similar groups of cells are compared first, reducing the chance of conflating noise with biological signal. In contrast, approaches that only assess groups of cells from a top-down perspective, pervasive in existing tools, risk masking signal from heterogeneous groups of cells, which can lead to inaccurate, underclustered results. Similarly, approaches that only assess groups of cells from a bottom-up perspective require the user to pre-determine where the bottom of the tree is, introducing the potential for underclustering as well.

When we identify clusters in single-cell data, we interpret those clusters as inherently useful cell groupings for subsequent analyses; therefore, the relatedness of clusters can be an informative metric. CHOIR preserves a record of all of the pairwise comparisons conducted before reaching the final set of clusters. This information can then be used to demonstrate the degree of relatedness of clusters or interrogate cell lineages (**Extended Data Fig. 1c**). In addition, CHOIR records feature importances found by the random forest classifier for each comparison, which represent the gene products or other features that best differentiate each cluster and can be used to aid cluster annotation. For instance, in the multi-omic human retina dataset, the feature importances measured in the comparison of CHOIR cluster 1, which represents rod cells, and CHOIR cluster 4, which represents cone cells, revealed features with differential gene expression and chromatin accessibility between these cell types (**Extended Data Fig. 1d–j**).

### Benchmarking results

Evaluating the performance of single-cell clustering algorithms presents challenges due to the lack of genuine “ground truth” information for most real single-cell sequencing datasets. Although simulated data provide a known ground truth and allow precise control of size and complexity, they cannot reproduce all aspects of real single-cell datasets. Furthermore, using published cell type annotations as a stand-in for ground truth cell identities risks biasing benchmarking results in favor of methods similar to those used in the original analysis. We therefore used complementary strategies in a three-tiered benchmarking approach to comprehensively compare the performance of CHOIR with that of 14 other unsupervised clustering methods^5,6,9,10,16–25^ (**Supplementary Table 3**) in the analysis of both simulated and real data (**Supplementary Fig. 1c** and **Supplementary Table 1**). We benchmarked the performance of each method across an array of parameter combinations, including the defaults, to assess the best possible performance in addition to the default performance. Overall, our results demonstrate that CHOIR outperforms available clustering methods across a range of species, cell numbers, tissue types, and sequencing modalities.

#### CHOIR accurately identifies simulated cell groups

We generated 100 simulated datasets representing 1, 5, 10, or 20 distinct ground-truth cell populations, ranging from 500 to 25,000 cells (**Fig. 2a**). Overall, CHOIR performed as well as or better than the 14 clustering methods used for comparison, as measured by the Adjusted Rand Index (ARI)^26^ and deviation from the ground-truth number of clusters (**Fig. 2b–i, Supplementary Fig. 2–21, Source Data 1**). Across the range of size and complexity represented by these simulated datasets, the computational time required by CHOIR scaled approximately linearly with the number of cells in each dataset (**Extended Data Fig. 2, Source Data 1**).

**Fig. 2:**
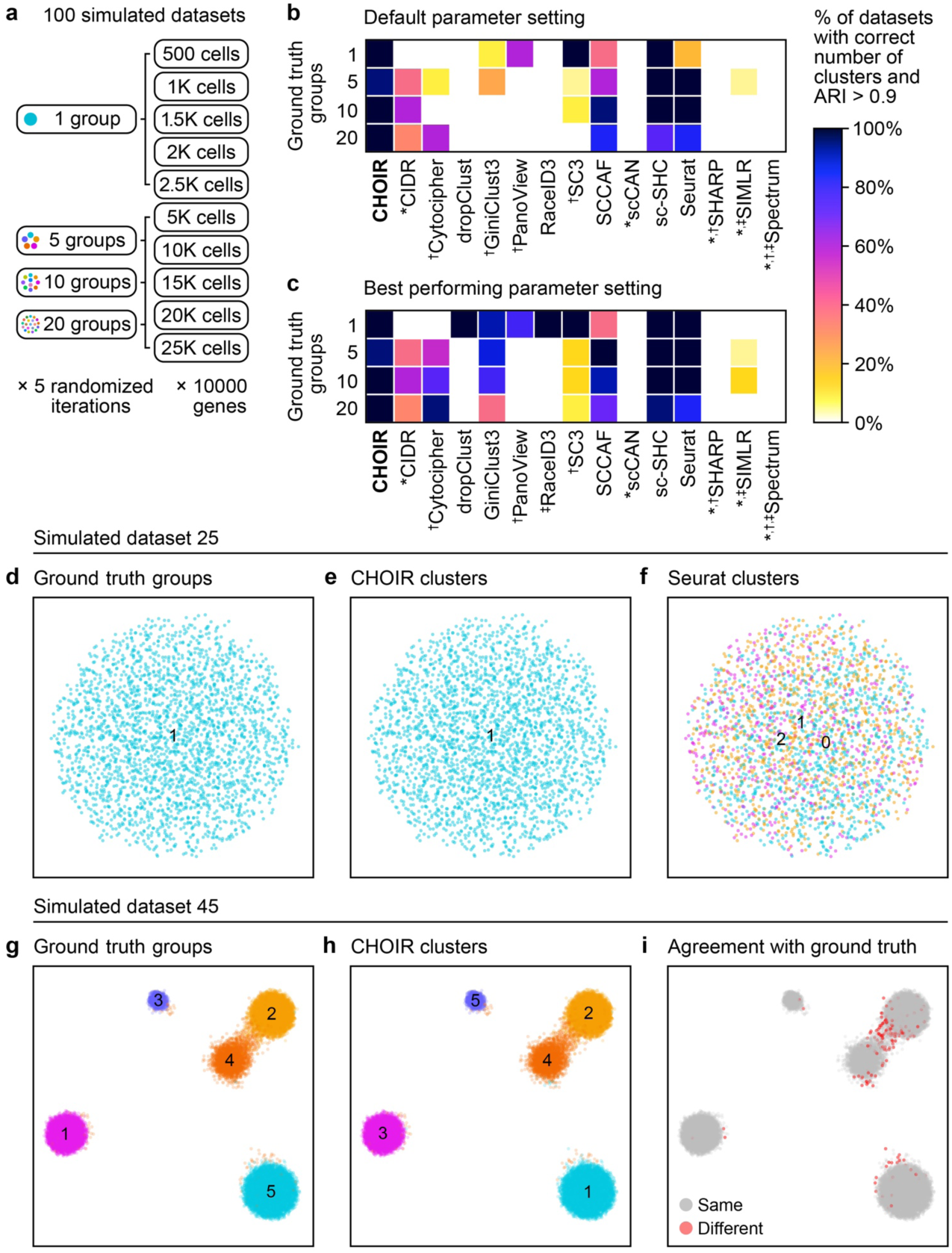
CHOIR outperforms 14 existing clustering methods across 100 simulated datasets. **a**. Schematic summarizing the 100 simulated datasets created using the R package Splatter. **b–c.** Heatmaps showing the percentage of simulated datasets in which the correct number of clusters was identified (all datasets) and with an ARI >0.9 (datasets with >1 ground truth groups) for the default parameter setting (**b**) or the best performing parameter setting (**c**) for each method. Symbols indicate clustering methods that *assumed ≥2 clusters, ^†^had some parameter settings that failed to run, or ^‡^had some parameter settings that did not complete within the maximum allotted runtime of 96 hours. For the CHOIR, CIDR, scCAN, SHARP, and Spectrum methods, the best-performing parameter setting in (**c**) was the same as the default parameter setting in (**b**). **d–f.** UMAP embeddings for simulated dataset 25, consisting of 2,500 cells and a single ground truth group of cells colored by the ground truth grouping (**d**) or the clusters achieved using the default parameters of CHOIR (**e**) or Seurat (**f**). **g–i.** UMAP embeddings for simulated dataset 45, consisting of 20,000 cells and five ground truth groups of cells colored by the ground truth groupings (**g**), the clusters achieved using the default parameters of CHOIR (**h**), and the agreement between these CHOIR clusters and the ground truth labels (**i**).

#### CHOIR avoids overclustering in simulated datasets consisting of a single population

Overclustering poses the risk of false discoveries of spurious new cell types or states. Within the set of 25 simulated scRNA-seq datasets representing a single ground truth cell population, CHOIR was one of only three methods (along with SC3 and sc-SHC) that consistently avoided overclustering (**Fig. 2a–f, Supplementary Fig. 2–6, Source Data 1**). Indeed, several of the clustering methods tested (CIDR, scCAN, SHARP, SIMLR, and Spectrum) assume a minimum of two clusters and never returned a single cluster. Even methods that permit the identification of a single cluster often misinterpret noise as structure, leading to overclustering. This weakness limits the utility of these methods for analyses where the presence or absence of discrete cell populations may be unknown, such as in subclustering analyses, analyses of cell lines, or analyses of homogeneous cell populations obtained by fluorescence-activated cell sorting (FACS). Although SC3 and sc-SHC consistently prevented overclustering across the simulated datasets consisting of a single ground truth population, these methods resulted in instances of underclustering across the more complex simulated datasets, in contrast to CHOIR (**Fig. 2b–c, Supplementary Fig. 7,12,17–19,21, Source Data 1**). The five existing methods with the best performance across these simulated datasets, Cytocipher, GiniClust3, SCCAF, sc-SHC, and Seurat, were used for comparison in the subsequent benchmarking assessments of CHOIR on real datasets. It is worth noting that many of the methods that performed poorly on these simulated datasets also required prohibitively long run times to benchmark on larger real datasets.

We also examined the performance of CHOIR by simulating the commonly used workflow in which clusters are manually selected for additional rounds of subclustering. Only CHOIR and sc-SHC consistently prevented overclustering when clusters identified in an initial round of clustering were subsetted and reclustered (**Supplementary Fig. 22, Source Data 1**). Thus, the clusters identified by CHOIR can be considered “terminal” clusters that do not require further rounds of subclustering.

#### CHOIR avoids underclustering in pools of distinct cell lines

To assess whether CHOIR also guards against underclustering, we evaluated the performance of CHOIR on a publicly available scRNA-seq dataset composed of pooled cancer cell lines^27^ (**Supplementary Table 1**). Although there might be distinct cell types or states within each cell line, distinct cell lines can be assumed to represent the minimal distinct identities in such a dataset. Therefore, multiple cell lines should never be grouped together into a single cluster. Across all parameter settings tested for the six clustering methods, only CHOIR was able to distinguish all 190 cell lines (**Fig. 3a–h, Supplementary Fig. 23, Supplementary Table 4**). CHOIR consistently prevented underclustering cells from distinct cell lines (**Fig. 3i**), as measured by the entropy of cluster accuracy (see **Methods**). The default parameter settings of Cytocipher, GiniClust3, SCCAF, sc-SHC, and Seurat resulted in the inclusion of multiple cell lines in a single cluster (**Figure 3d–h, Supplementary Fig. 23e–i**), providing clear examples of underclustering.

**Fig. 3:**
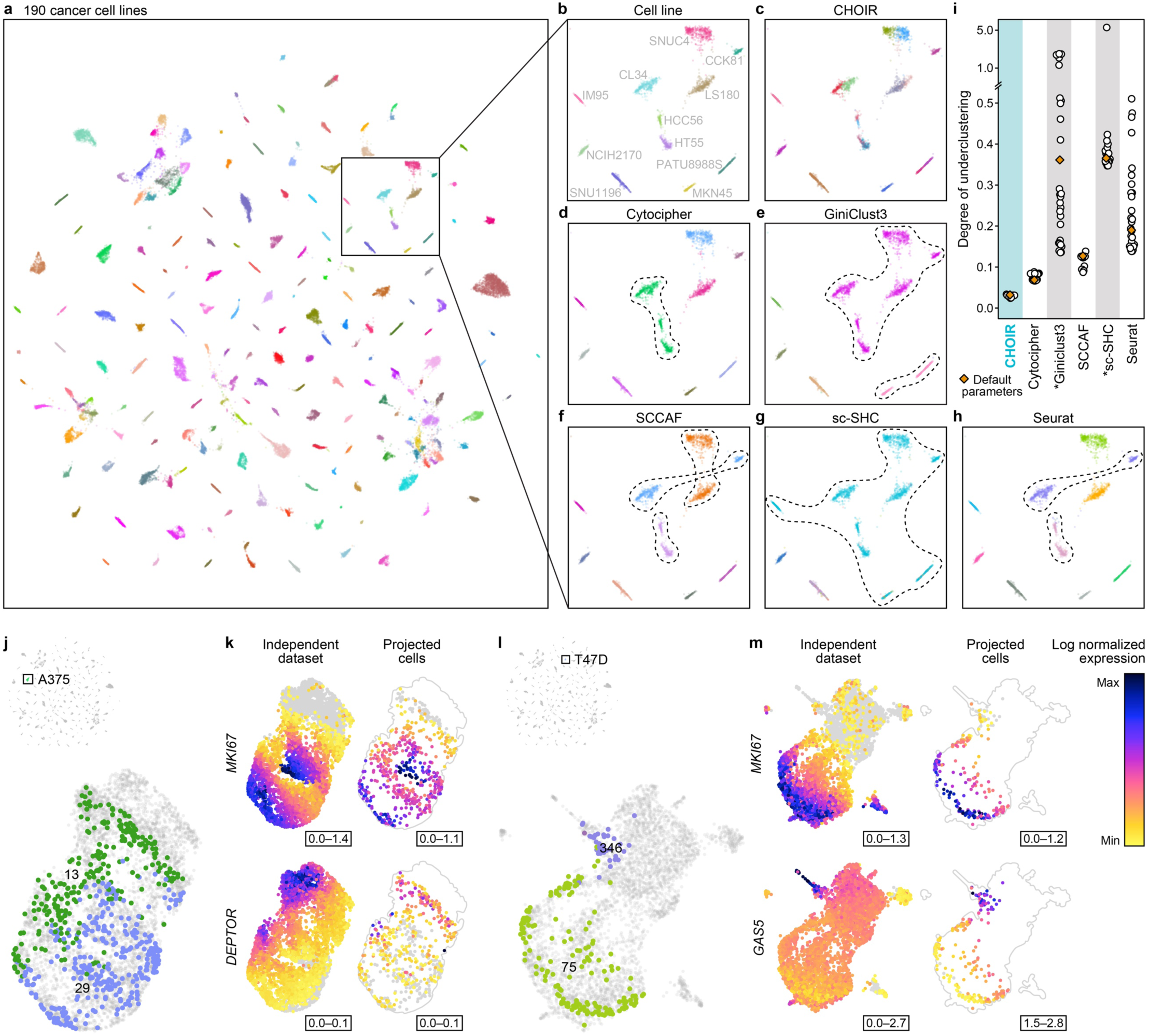
CHOIR prevents underclustering in an scRNA-seq dataset consisting of pooled cell lines. **a**. UMAP embedding of a pooled cancer cell line scRNA-seq dataset from Kinker et al. 2020 consisting of 48,879 cells, colored according to the 190 cancer cell lines. **b–h.** UMAP embedding showing the cells within the highlighted square in (**a**), colored by cell line (**b**) and the clusters identified by CHOIR (**c**), Cytocipher (**d**), GiniClust3 (**e**), SCCAF (**f**), sc-SHC (**g**), and Seurat (**h**), using the default parameters for each method. Dashed lines indicate instances where the clustering method failed to distinguish individual cell lines, resulting in multiple cell lines grouped within a single cluster. **i**. The entropy of cluster accuracy for all method and parameter combinations tested, representing the degree of underclustering. *Some parameters tested for GiniClust3 and sc-SHC resulted in a single cluster, for which the entropy of cluster accuracy could not be computed. **j**. UMAP embedding showing CHOIR cluster 13 and cluster 29 identified within the A375 cell line projected onto the dimensionality reduction space of an independent scRNA-seq dataset consisting of 4,794 A375 cells. **k**. Expression levels of proliferation marker *MKI67* and tumor suppressor *DEPTOR* across the A375 cells from the Yang et al. 2021 independent dataset (left) and the projected cells from Kinker et al 2020 (right). **l**. UMAP embedding showing CHOIR cluster 75 and cluster 346 identified within the T47D cell line projected onto the dimensionality reduction space of an independent scRNA-seq dataset consisting of 4,582 T47D cells. **m**. Expression levels of proliferation marker *MKI67* and growth arrest marker *GAS5* across the T47D cells from the Dave et al. 2023 independent dataset (left) and the projected cells from Kinker et al 2020 (right).

Within some cell lines, CHOIR distinguished multiple clusters, raising the question of potential overclustering if those clusters failed to represent distinct cell states with relevant marker genes. For two such cell lines (A375 melanoma cells and T47D breast cancer cells), we therefore projected the clusters identified by CHOIR onto larger, independent datasets from the same lines^28,29^ (**Supplementary Table 1**) to confirm whether these clusters represent biologically distinct cell populations. In both cases, the clusters identified by CHOIR projected onto distinct populations of cells within the larger independent datasets (**Fig. 3j–m**). For both cell lines, the two clusters identified by CHOIR within each cell line represented proliferating and non-proliferating cells. The proliferating cells expressed high levels of proliferation markers *MKI67* and *PCNA*^30,31^, whereas the non-proliferating cells expressed high levels of *DEPTOR*, an mTOR inhibitor and tumor suppressor^32^, and *GAS5*, a lncRNA known to be enriched during cell cycle arrest^33^ (**Fig. 3k,m**). These results confirm that the subclusters identified by CHOIR within a single cell line indeed represent distinct cell states and also highlight the ability of CHOIR to identify such states from relatively few total cells.

#### Orthogonal validation of CHOIR using multi-modal datasets

Multi-modal datasets provide a useful opportunity to leverage orthogonal data from the same cells to assess whether clusters are biologically distinct. For this purpose, we performed clustering on one modality and assessed the resulting clusters for differential features in the other modality. We collected two publicly available datasets with different types of multi-modal data (**Supplementary Table 1**): (1) CITE-seq data from human peripheral blood mononuclear cells (PBMCs) encompassing both scRNA-seq and antibody-based surface marker proteomics^34^ and (2) sci-Space data from developing mouse embryos comprising snRNA-seq data and spatial coordinates^35^. For each dataset, we benchmarked the performance of CHOIR against the five best-performing methods prioritized above (Cytocipher, GiniClust3, SCCAF, sc-SHC, and Seurat).

In the multi-omic CITE-seq dataset containing snRNA-seq and surface marker protein data from the same 159,827 human PBMCs^34^, each clustering method was applied only to the RNA modality. The surface marker protein data, comprising 228 oligonucleotide-conjugated antibodies, was then used to assess the minimum number of differentially expressed proteins among the 50 nearest cluster pairs identified by each of the clustering algorithms. We then interpreted observation of a pair of clusters with no differentially expressed proteins between them to be indicative of overclustering, as the ideal method would identify the highest number of clusters while still maintaining some differentially expressed proteins between every cluster pair. Of the six clustering methods tested, only CHOIR maximized the number of clusters without potentially overclustering the data, as evidenced by the fact that each cluster differed from its neighboring clusters in the expression of multiple proteins (**Fig. 4a– c, Supplementary Fig. 24a–f, Supplementary Table 5**). However, we note that the total number of features used for clustering from the RNA modality was much larger than the total number of features used for differential expression analysis from the protein modality. We therefore refer to the RNA-based distinction of clusters that did not differ by protein detection as “likely overclustering.”

**Fig. 4:**
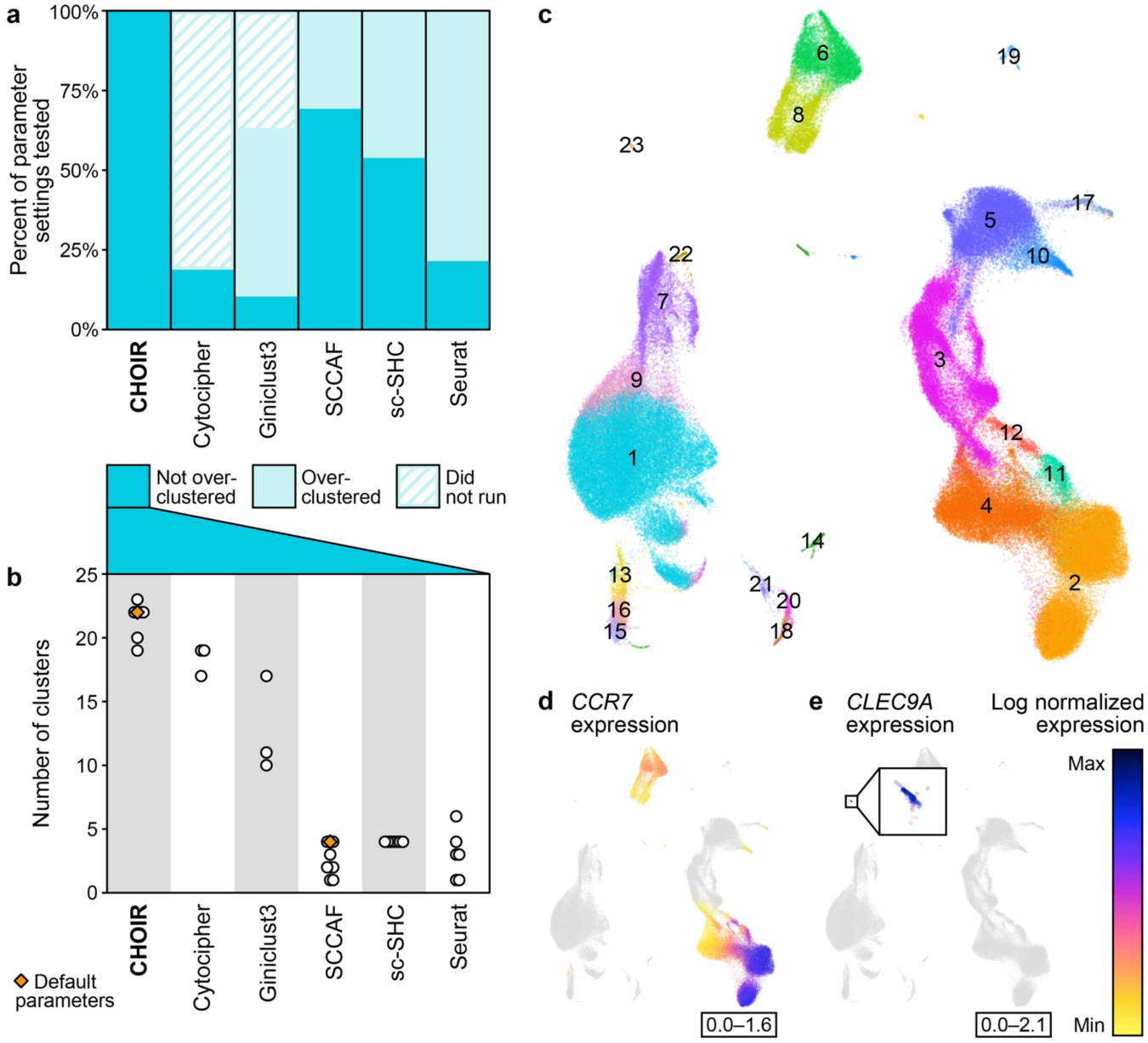
Multi-omic data enables orthogonal validation of identified clusters. **a**. A stacked barplot showing the percentage of parameter settings for each method tested on the Hao et al. 2021 human PBMC CITE-seq dataset that did (not overclustered) or did not (likely overclustered) result in at least one differentially expressed protein in all 50 closest pairwise cluster comparisons, or that failed to run. **b.** A dot plot showing the number of clusters that was identified by the subset of all method and parameter combinations tested that did not result in likely overclustering (**a**). **c.** UMAP embedding of cells from the Hao et al. 2021 human PBMC CITE-seq dataset colored according to the 23 clusters identified by the parameter settings for CHOIR that maximized the number of clusters while preventing likely overclustering. **d–e.** UMAP embedding as in panel (**c**), but colored according to the expression level of naïve T cell marker *CCR7* (**d**) or the expression level of conventional dendritic cell type 1 marker *CLEC9A* (**e**). Because of the small size of Cluster 23, a zoom-in window was used to display *CLEC9A* expression in panel (**e**).

Although all parameter settings tested for Cytocipher avoided likely overclustering, as indicated by differentially expressed proteins between the clusters, all of these parameter settings resulted in fewer clusters than those obtained with CHOIR, indicating possible underclustering by Cytocipher. Supporting this notion, CHOIR identified a population of *CCR7*-expressing naïve T cells, a subset of T lymphocytes that are antigen-inexperienced^36^ (**Fig. 4d, Supplementary Fig. 24a–b, Supplementary Table 5**), and a population of *CLEC9A*-expressing conventional type 1 dendritic cells, a small population of antigen-presenting cells that are key to generating cytotoxic T cell responses^37^ (**Fig. 4e, Supplementary Fig. 24a–b, Supplementary Table 5**), which were not identified in any of the parameter settings tested in Cytocipher. Although some parameter settings for GiniClust3, SCCAF, and Seurat yielded more clusters than CHOIR, in all such cases, pairwise comparisons of these clusters revealed no differentially expressed proteins, implying likely overclustering (**Fig. 4a–b**). No parameter settings tested for sc-SHC identified as many clusters as were identified by CHOIR, and the comparison of some clusters obtained with sc-SHC showed no differentially expressed proteins, implying both under– and overclustering. Furthermore, although Harmony batch correction was applied during CHOIR, Cytocipher, SCCAF, and Seurat clustering, only CHOIR and Cytocipher avoided the spurious segregation of clusters of the same cell type based on batch effects, whereas GiniClust3, Seurat and SCCAF were susceptible to this confound (**Supplementary Fig. 24g–j**).

In the sci-Space dataset^35^, which consisted of sagittal sections of whole E14 mouse embryos, the clustering methods were again applied only to the RNA modality, without reference to the spatial coordinate information. Using the default parameter settings of CHOIR, 42 clusters were identified (**Fig. 5a, Supplementary Table 6**). Given that the E14 developmental period is characterized by the formation of many organ systems, we would expect to identify not only anatomically dispersed cell types, but also a number of organ-specific clusters that are becoming spatially constrained (**Fig. 5b–c**). Indeed, some cell types, for example, keratinized epithelial cells, were found to be distributed throughout the body (**Fig. 5d**), whereas other cell types were found to be restricted to circumscribed anatomical locations, for example, hepatocytes in the liver (**Fig. 5e**) and mesenchymal cells in the lung (**Fig. 5f**). We used such differences in distribution as an external validation of the biological relevance of clusters identified. For each tissue section, we evaluated the percentage of all clusters that had a highly localized spatial distribution, defined as a mean radial area that was less than 5% of the approximate whole embryo section area. Across all parameter settings tested for each method, CHOIR obtained the highest mean percentage of spatially localized clusters (**Fig. 5g**). The spatial restriction of these clusters to anatomically defined regions suggests that these clusters represent biologically distinct cell populations, a conclusion that was further validated by the expression of known developmental marker genes, as delineated below.

**Fig. 5:**
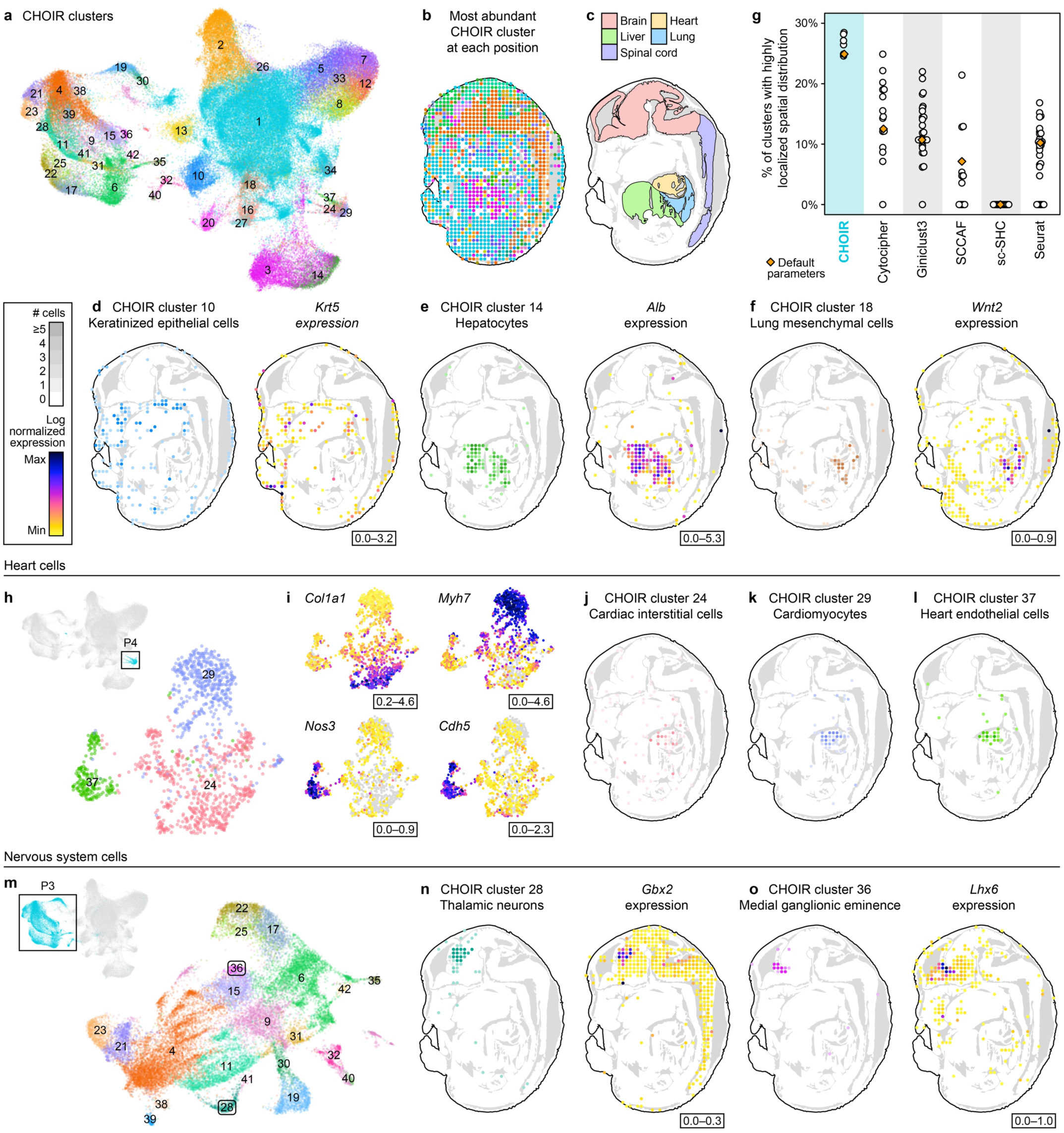
CHOIR identifies anatomically localized clusters in a spatially resolved snRNA-seq dataset. **a**. UMAP embedding colored according to the 42 clusters identified by applying the default parameters of CHOIR to the snRNA-seq features of the Srivatsan et al. 2021 whole mouse embryo sci-Space dataset. Cluster colors persist throughout this figure. **b.** CHOIR cluster with the highest number of cells at each spatial coordinate position mapped onto a single section. Slide 14 is shown throughout this figure because it had the highest number of recovered nuclei and positions. **c.** Anatomical boundaries, adapted from annotations used in Srivatsan et al. 2021. **d–f.** Cluster distributions and corresponding marker gene expression levels for CHOIR cluster 10, keratinized epithelium cells (**d**); cluster 14, hepatocytes (**e**); and cluster 18, lung mesenchyme cells (**f**) mapped onto spatial coordinates. For plots showing CHOIR clusters, the darker the shade of any given color, the more cells of the respective cluster were detected at the indicated location. **g.** Mean percentage of all clusters per section that had a highly localized spatial distribution for all method and parameter combinations tested (see **Methods**). For Cytocipher, some parameter settings failed to run. **h–i.** UMAP embedding computed on the subsetted dimensionality reduction of parent cluster P4 identified using default parameters of CHOIR colored by CHOIR cluster (**h**) and the expression levels of connective tissue marker gene *Col1a1*, cardiomyocyte marker gene *Myh7*, and endothelial cell marker genes *Nos3* and *Cdh5* (**i**). See inset legend next to panel (**d**). **j–l.** Cluster distributions for CHOIR cluster 24, *Col1a1*-expressing cardiac interstitial cells (**j**), cluster 29, *Myh7*-expressing cardiomyocytes (**k**), and cluster 37, *Nos3*/*Cdh5*-expressing heart endothelial cells (**l**) mapped onto spatial coordinates. The darker the shade of any given color, the more cells of the respective cluster were detected at the indicated location. See inset legend next to panel (**d**). **m.** UMAP embedding computed on the subsetted dimensionality reduction of parent cluster P3 consisting of nervous system cells colored by the clusters identified by CHOIR using the default parameters. **n–o.** Cluster distributions and corresponding marker gene expression levels for CHOIR cluster 28, *Gbx2*-expressing thalamic neurons (**n**), and cluster 36, *Lhx6*-expressing medial ganglionic eminence interneuron progenitors (**o**). For plots showing CHOIR clusters, the darker the shade of any given color, the more cells of the respective cluster were detected at the indicated location. See inset legend next to panel (**d**).

CHOIR identified several clusters with highly specific anatomical localization that were missed by other clustering methods. For example, CHOIR was the only method that distinguished three distinct cell types of the heart: cardiac interstitial cells, cardiomyocytes, and heart endothelial cells (**Fig. 5h–l, Supplementary Fig. 25**). The cardiac interstitial cell cluster, the cardiomyocyte cluster, and the heart endothelial cell cluster identified by CHOIR were all anatomically localized to the heart across all sections (**Fig. 5j–l**). However, these cell clusters were appropriately differentiated based on the expression of marker genes for connective tissue (*Col1a1*, *Col14a1*^38^), endothelial cells (*Cdh5*, *Nos3*, *Pecam1*, *Vwf*^39,40^), or cardiomyocytes (*Myh6*, *Myh7*^41^), respectively, and were clearly distinct when viewed in a refined dimensionality reduction (**Fig. 5h–i, Supplementary Fig. 25**).

CHOIR was also the only method whose default parameters distinguished certain anatomically restricted neuronal subpopulations. In total, CHOIR identified 22 nervous system clusters, ranging in size from 37 to 6,875 cells (**Fig. 5m**). Of these, Cluster 28 was characterized by high *Gbx2* expression and represents a population of neurons that are localized within the developing thalamus^42^ (**Fig. 5n, Extended Data Fig. 3**). Cluster 36 represents the pool of medial ganglionic eminence interneuron progenitors characterized by high expression of *Lhx6*^43^ (**Fig. 5o, Supplementary Fig. 26**). Additionally, CHOIR identified a neuronal progenitor pool of the spatially restricted E14 dorsal pallium characterized by expression of *Dbx1* and *Neurog1*^44–46^ (**Supplementary Fig. 27**), and a cerebellar population of neurons with characteristic *Fgf3* expression^47^ (**Supplementary Fig. 28**).

Across all parameter settings tested for each method, CHOIR was the only method to identify all of the above-mentioned heart and brain cell types, in addition to many other examples (**Supplementary Fig. 25–29, Supplementary Table 6**). Some non-default parameters for GiniClust3 and Seurat also identified several of these cell types; however, those parameter settings resulted in severe overclustering of other cell types (**Supplementary Fig. 29**).

Thus, using a tiered benchmarking approach that leveraged both simulated and real datasets (**Supplementary Fig. 1c**), we found that CHOIR successfully prevented both underclustering and overclustering. CHOIR accurately identified simulated ground truth cell groups, avoided overclustering of simulated datasets composed of a single ground truth population, and avoided underclustering of pooled cancer cell lines. Using multi-omic datasets, we demonstrated that the clusters identified by CHOIR are clearly distinct in an orthogonal modality, indicating that CHOIR is able to detect biologically relevant clusters that are difficult or impossible to reliably identify by other clustering methods.

## DISCUSSION

Defining reliable clusters that represent distinct cell types and states is critical to gaining biological insights from single-cell analyses. However, the subjective nature of commonly used approaches to clustering does not sufficiently protect against overclustering or underclustering of the data, which can lead to misleading conclusions about fundamental and disease-relevant biological processes. In addition, many existing clustering approaches preferentially identify clusters of similar sizes, often resulting in overclustering of common homogeneous cell types and underclustering of rare cell populations. CHOIR addresses these challenges by incorporating statistical inference into the clustering process, reducing the chance of spurious discoveries. By tying cluster identification to a significance threshold, CHOIR ensures that the clusters identified all meet a standardized criterion. Common homogeneous cell populations are not spuriously split due to noise, and rare populations are identified as distinct even when the total number of cells is low. The primary tradeoff when using a clustering tool such as CHOIR is the increased processing time required. For large datasets, CHOIR could be orders of magnitude slower than a single iteration of standard Louvain or Leiden clustering. However, although CHOIR has a longer run time than standard Louvain or Leiden clustering, it saves many hours of manual optimization of cluster definition and only needs to be performed once per dataset, as the clusters identified by CHOIR do not require further rounds of subclustering. Moreover, in the course of the hierarchical clustering procedure, CHOIR can define marker features for each cluster, further assisting in downstream cluster annotation.

Existing hierarchical clustering approaches exclusively utilize either a top-down or bottom-up approach. Top-down approaches start with all cells in a single group and proceed to attempt to iteratively divide that group until further division is deemed unnecessary. While this is intuitive, it risks masking signal from highly variable groups of cells, sometimes leading to underclustering (**Extended Data Fig. 1c**). Bottom-up approaches avoid this pitfall by ensuring that the most similar groups of cells are compared first, reducing the chance of conflating noise with heterogeneous biological signal. However, it is challenging to know what the true “bottom” of the tree is without first constructing the tree from the top down. CHOIR leverages the strengths of both approaches and is implemented in two main steps: a top-down construction of a hierarchical clustering tree, followed by a bottom-up pruning of the branches of that clustering tree. These two steps are used to prevent underclustering and overclustering, respectively. To our knowledge, CHOIR is the only available method to take this two-step approach. Although Cytocipher and SCCAF also implement a bottom-up approach to prevent instances of overclustering, they rely on the user to determine the input Louvain or Leiden resolution, subjectively setting a threshold at which the dataset is deemed to be overclustered. These methods also do not assess whether underclustering has occurred and are, thus, vulnerable to it. In contrast, sc-SHC takes only a top-down approach. This method obviates the need for the user to determine the input clusters but is still susceptible to underclustering, due to the top-down approach itself.

In addition to combining top-down and bottom-up steps, CHOIR implements an additional strategy to prevent underclustering: as the initial hierarchical clustering tree is constructed, new dimensionality reductions and sets of highly variable features are generated for a preliminary set of high-level clusters that typically represent the major cell classes present in the dataset. For random forest classifier comparisons applied to clusters that stem from the same parent cluster, the subsetted dimensionality reduction and feature set are used. This allows CHOIR to identify more nuanced differences between cell subtypes or states that are not easily detected using existing clustering methods, as highlighted by the identification of proliferation states within cancer cell lines by CHOIR (**Fig. 3j–m**), as well as the identification of functionally distinct cell types in the heart (**Fig. 5h–l**) and cell types specific to different brain regions (**Fig. 5m–o**) within the challenging, more continuous data structure observed in the dataset from developing mouse embryos.

To achieve accurate clustering results, these nuanced differences between cell subtypes or states must not be conflated with batch-driven noise—a large potential source of overclustering. Other methods such as Cytocipher and SCCAF purport to prevent overclustering, but neither algorithm has batch correction checks built into the clustering method itself. Therefore, although the initial clusters may be determined using a batch-corrected dimensionality reduction, the cell-by-gene count matrix itself would have to be batch-corrected to prevent batch effects from biasing the ultimate cluster labels that pass the significance and accuracy thresholds imposed, and this is not routinely done in single-cell analyses. CHOIR offers a unique solution to this challenge. For datasets that require batch correction, CHOIR uses an internal Harmony-based batch correction approach^48^. Harmony is applied to generate the dimensionality reductions used, and batch labels are additionally used to subset the input to the random forest classifiers, so that for each cluster comparison only cells from the same batch are compared. In instances where the two clusters being compared have very few cells belonging to the same batch, these clusters are assumed to be batch-confounded instances of the same cell population and are merged. For example, if cluster 1 predominantly consists of cells belonging to batch A, but cluster 2 predominantly consists of cells belonging to batch B, these two clusters would be merged. Thus, batch information is used at every stage of the clustering algorithm in order to avoid overclustering due to batch confounds.

Beyond confounds due to batch effects, factors such as read depth and cell count will always affect the granularity with which cell types and states can be distinguished. Even high-quality clustering results are simply proxies for the highly continuous landscape of real biological features. Although CHOIR performs well on datasets across a range of developmental continuity, there may be instances where trajectory analyses should be combined with clustering approaches in order to represent the nuances of developmental lineages. Additionally, as cell counts in typical single-cell experiments increase by orders of magnitude due to the increasing accessibility of sequencing technologies, new methods, such as meta-cells^49,50^ or ‘sketching’^10,51^, may be needed to maintain the same computational efficiency. As the size and diversity of available single-cell datasets increase, so too will the absolute number of bona fide cellular states. Although CHOIR does not define ‘singleton’ clusters composed of a single cell, it is typically able to distinguish rare clusters with as few as 20–30 cells and clusters of cells that make up less than 0.05% percent of a dataset, highlighting an enabling strength of this new algorithm.

In light of the ever-widening field of single-cell sequencing technologies, we extended the utility of CHOIR beyond scRNA-seq data by implementing computational methods free of assumptions about the distribution of the data. The random forest classifier and permutation test framework used by CHOIR were specifically chosen because they are free of such assumptions. CHOIR is therefore amenable to feature matrices of any modality and is applicable to multi-modal data with an arbitrary number of features and modalities. Consequently, CHOIR is uniquely poised to address both current needs and future developments in single-cell analysis.

Notably, across an array of dataset sizes, modalities, and biological contexts, the default parameters implemented in CHOIR were consistently either the optimal parameters or yielded similar results as the optimal parameter settings, a feature that enhances performance and mitigates subjectivity in cluster identification. CHOIR was able to identify unique biologically relevant cell groups in challenging contexts such as within clonal cell lines or during mouse development. In light of the strengths described above, this computational approach could help standardize the process of cell type and cell state identification across the diversity of modalities and biological contexts of interest to the single-cell research community.

## METHODS

### Details of the CHOIR algorithm

#### Random forest classifiers used in a permutation test framework

CHOIR operates around a central algorithmic cog: a permutation test using random forest classifiers to learn and predict the cluster labels of cells in a pair of clusters. The cells belonging to the pair of clusters are split equally into a training set and test set. If the two clusters do not contain the same number of cells, the larger cluster is downsampled to create a balanced representation of the two clusters in the training and test set. A random forest classifier is then trained using a cell by feature matrix. By default, CHOIR will use only the highly variable features. For multi-omic datasets, highly variable features from all modalities are used jointly as the input for the random forest classifier. Subsequently, the trained random forest classifier is used to predict the cluster assignments of the cells in the test set. This process yields a prediction accuracy score between zero and one, where one represents perfect prediction. Higher prediction scores indicate that the two clusters are more different, and thus more readily distinguishable. In parallel, CHOIR shuffles the cluster labels and repeats the same process using both true and permuted labels on bootstrapped samples over a number of iterations (100 by default), resulting in a permutation test that compares the true prediction accuracy for the clusters to the prediction accuracy for a chance division of the cells into two random groups. This permutation testing yields a p-value that determines whether the two clusters are merged or kept separate. The significance threshold used can be adjusted according to an alpha parameter. The default significance level used by CHOIR is 0.05 with Bonferroni multiple comparison correction.

For datasets that require batch correction, CHOIR uses a batch-aware approach to run the permutation tests. When batch correction is enabled, only cells from the same batch are compared in a single random forest classifier iteration. The batches present in the data alternate across all of the iterations of the permutation test. In addition, if the two clusters being compared have very few cells belonging to the same batch, they are merged to avoid batch confounds. This batch correction approach assumes that any experimental groups of interest that may have unique cell types or states are not batch-confounded.

Because the iterations of the permutation test are parallelized, the efficiency of CHOIR greatly improves with additional computational cores. In addition, CHOIR applies three filters that reduce the total number of comparisons performed. First, CHOIR imposes a minimum accuracy threshold for the random forest classifier predictions, below which clusters will be automatically merged. This accuracy threshold defaults to random chance (0.5); higher values can be set in order to identify a more conservative set of clusters. Second, by default, CHOIR only runs the permutation test comparison if the two clusters share at least one nearest neighbor. Clusters with no nearest neighbors remain separate. The minimum number of nearest neighbors required can be adjusted by the user. Third, CHOIR records the distance (in the generated dimensionality reduction space) between each cluster and its nearest distinguishable neighbor. Multiplied by a set value, this distance is used to generate a distance threshold above which this cluster is not merged with any other clusters. By default, this threshold is set at a two-fold increase in distance. CHOIR also uses downsampling to increase efficiency for larger datasets. By default, downsampling occurs at each random forest classifier comparison. The downsampling rate is determined based on the overall dataset size or can be set by the user.

#### Generating the initial hierarchical clustering tree

The first step in the CHOIR algorithm is the top-down generation of a hierarchical clustering tree. In the first layer of this clustering tree, all cells belong to a single cluster. The first section of the tree is referred to as the “root tree.” Using all cells, CHOIR identifies a set of features with highly variable levels of expression from the provided cell by feature matrix. By default, CHOIR uses 25,000 variable features for ATAC-seq data and 2,000 variable features for other types of data. CHOIR then computes a dimensionality reduction for all cells, using one of three methods: principal component analysis (PCA), latent semantic indexing (LSI), or iterative LSI. PCA is used as the default, except for ATAC-seq data. For datasets that require batch correction, CHOIR generates a Harmony-corrected dimensionality reduction^48^. For multi-omic datasets, a joint dimensionality reduction is created across all of the provided modalities. CHOIR then computes a nearest neighbor adjacency matrix. To generate the layers of the clustering tree, CHOIR uses either Louvain^2^ or Leiden^3^ clustering with increasing resolution parameters. CHOIR incrementally increases the resolution parameter at each level of the clustering tree. Because increases in the resolution parameter for Louvain and Leiden clustering do not lead to strictly hierarchical divisions of clusters, CHOIR uses the tool MRtree^52^ to reshape clustering trees into a strictly hierarchical clustering tree.

At each level of the emerging root tree, the silhouette score is assessed. By default, an approximate silhouette score is used to avoid generating full distance matrices. Subdivision of the root tree continues until the silhouette score has reached a maximum score that is not exceeded in the subsequent levels of the tree. The root tree is then terminated at the level with the maximum silhouette score. This level is referred to as the “parent clusters.” Each parent cluster is then subsetted, a new set of highly variable features is identified, and a new dimensionality reduction and nearest neighbor adjacency matrix are generated. These new elements are used as the basis for a “subtree” originating from the parent cluster. Each of the resulting subtrees continues to be subdivided until the farthest pair of nearest neighboring clusters is found to be overclustered by the permutation test approach. At this level, the clustering tree is terminated, as it is likely that most, if not all, of the clusters are overclustered.

#### Pruning the hierarchical clustering tree

To prune back the branches of the hierarchical clustering tree, CHOIR iterates through each branch point of the clustering tree using a bottom-up approach. At each branch point, pairs of clusters are compared with the permutation test approach. For these permutation tests, the input matrices used for the random forest classifiers depend on the parent cluster origins of the two clusters being compared. If both clusters are from the same subtree, the highly variable features, dimensionality reduction, and nearest neighbor adjacency matrix generated for that subtree are used as input to the permutation test framework. Otherwise, the root tree data is used.

When all levels of the clustering tree have been assessed, the resulting clusters can be assumed to have no instances of overclustering. If any of the resulting clusters are identical to clusters in the bottom tier of the tree (where pruning began), these clusters are subdivided and undergo additional rounds of pruning, ensuring that no instances of underclustering remain. Thus, CHOIR assesses both overclustering and underclustering to identify the final clusters.

For users who already have a set of clusters generated by a different tool and wish to assess whether these clusters are under-or overclustered, CHOIR can accept an externally generated clustering tree, dimensionality reduction, and nearest neighbor adjacency matrix, and then proceed in the same manner as described above.

### Simulated data

Synthetic datasets were generated using the R package Splatter (1.20.0)^53^, which simulates scRNA-seq data from either single or multiple cell populations based on a gamma-Poisson distribution. We generated a set of 100 simulated datasets, spanning 1 to 20 ground truth groups and 250 to 20,000 cells, with 10,000 features each (**Fig. 2a**). Each group and cell number combination was repeated 5 times, using randomized parameter values.

### Benchmarking

#### Pre-processing

For each dataset, a standardized pre-processing pipeline was applied using Snakemake (7.32.4)^54^. Depending on the file format of the input files (bam, fastq, or mtx) available for each dataset, Cell Ranger (7.2.0) and Cell Ranger ARC (2.0.1) functions were applied to generate a count matrix. For data available as bam or fastq files, analyses of human and mouse data were performed using the hg38 genome and mm10 genome, respectively, using refgenie^55,56^ for genome resource management.

For sn/scRNA-seq data, a series of quality control steps were applied: (1) Cell Ranger or emptyDrops (DropletUtils 1.16.0)^57^ was used to identify droplets containing only ambient RNA, (2) the number of reads per cell, number of features per cell, and percent mitochondrial reads were used to filter out low-quality cells or nuclei that were more than three median absolute deviations from the median^58^, and (3) doublets were detected with DoubletFinder (2.0.3)^59^ using default parameters. Data were supplied to clustering methods as raw counts or log normalized counts. For CHOIR and Seurat, datasets larger than 25,000 cells were provided as BPCells (0.1.0) counts matrices for maximal efficiency.

For snATAC-seq data, a series of quality control steps was applied using ArchR (1.0.3)^12^. Cells with transcription start site (TSS) enrichment scores <2 or fragment counts <2500 were excluded. ArchR was additionally used to detect and filter out doublets. For the Wang et al. 2022^15^ multi-omic ATAC-seq and RNA-seq dataset, clustering was conducted on the shared set of cells that passed quality control thresholds in both pre-processing pipelines.

#### Clustering

Each benchmarking dataset was clustered using CHOIR alongside a set of existing methods (**Supplementary Table 3**) according to the data type, across a range of reasonable parameter measures informed by available method documentation, including the default parameter values for each method. For the simulated datasets, 14 existing clustering methods were used for comparison. Of these, the five methods with the best performance, Cytocipher, GiniClust3, SCCAF, sc-SHC, and Seurat, were benchmarked using the real single-cell datasets. For datasets that require batch correction, CHOIR uses an internal Harmony-based batch correction approach^48^. Harmony batch correction was applied to the other methods as well, with the exception of sc-SHC, which does not use a dimensionality reduction for the input to clustering and has an internal batch-correction method, and GiniClust3, which is not compatible with batch correction. All clustering methods were run using 16 cores (if parallelized). Computational time and memory usage were recorded, and a maximum runtime of 96 hours was imposed. CHOIR numbers clusters in order from largest to smallest. A standard color palette for the cluster order was used throughout the analyses shown. For methods in which the resulting clusters were not numbered from largest to smallest, the color palette was still applied in order from largest to smallest, to aide visual comparison.

#### Performance Metrics

For simulated datasets, we were primarily concerned with the similarity between the generated clusters and the ground truth groups, which was measured using the ARI^26^ for simulated datasets with more than one ground truth group. For the cancer cell line dataset, we assumed that the cell line identities represented the minimal distinct identities present in the dataset. We quantitatively assessed the degree of underclustering for each method by using the entropy of cluster accuracy^60^ to measure the diversity of cell lines present in each cluster. For the entropy of cluster accuracy, higher values indicate that multiple cell lines are grouped in a single cluster, and lower values indicate a prevention of underclustering.

For the multi-omic CITE-seq dataset, clusters were computed on the RNA-seq modality, and separation along the orthogonal surface marker protein modality was evaluated. For these clusters, we identified the closest 50 cluster pairs as measured by the centroid distance in the reduced PCA dimensions generated using the default parameters of CHOIR. Using edgeR (3.36.0)^61^ within the Libra (1.0.0)^62^ package interface, we then conducted pairwise differential expression analysis of the surface marker protein data, using pseudobulked values for each sample for each of the cluster pairs. The minimum number of differentially expressed genes with an FDR-adjusted p-value <0.05 was assessed, as well as the number of clusters obtained. These metrics were used to identify methods that maximize the number of clusters while preserving pairwise differences among clusters. Because the surface marker protein data was not used to assign the cluster labels, it represents an external validation of the cluster differences. For the sci-Space dataset, clusters were also computed using the snRNA-seq data, without any use of the spatial coordinate data; the cluster labels were then imposed upon the respective spatial coordinates of each section to assess the spatial distribution of clusters. For each section, the mean percentage of all clusters that had a highly localized spatial distribution across the section, defined as a mean radial area <5% of the approximate whole embryo section area, was computed for all parameter settings tested for each clustering method. A rectangular area was computed as an approximation of the sci-Space sampling area for each section. Expression levels of corresponding marker genes for each cluster were visualized on Uniform Manifold Approximation and Projection (UMAP) embeddings and were also mapped onto spatial coordinates. MAGIC (2.0.3)^63^ was used to impute expression levels for visualizations only.

## DATA AVAILABILITY

All datasets were acquired from the accession numbers provided in the original publications (**Supplementary Table 1**). The simulated datasets generated here, as well as other analysis files, are available on Zenodo at https://doi.org/10.5281/zenodo.14641222.

## CODE AVAILABILITY

CHOIR is available as an R package at https://github.com/corceslab/CHOIR, and documentation for CHOIR is hosted at https://www.CHOIRclustering.com. The code to reproduce the analyses from this paper is available at https://github.com/corceslab/2025_Sant_CHOIR.

## ACKNOWLEDGEMENTS

We thank members of the Corces and Mucke laboratories for helpful comments, and A. Dalton and R. Mott for administrative assistance. This work was supported by National Institutes of Health grants P01 AG073082 (L.M. and M.R.C.) and U01 AG072573 (M.R.C.). C.S. was supported by a National Science Foundation Graduate Research Fellowship.

## CONTRIBUTIONS

C.S., L.M., and M.R.C. conceived the project. C.S. led the design of the CHOIR software with input from L.M. and M.R.C. C.S. led the single-cell analyses presented in this paper. C.S., L.M., and M.R.C. wrote the manuscript.

## EXTENDED DATA FIGURES

**Extended Data Fig. 1:**
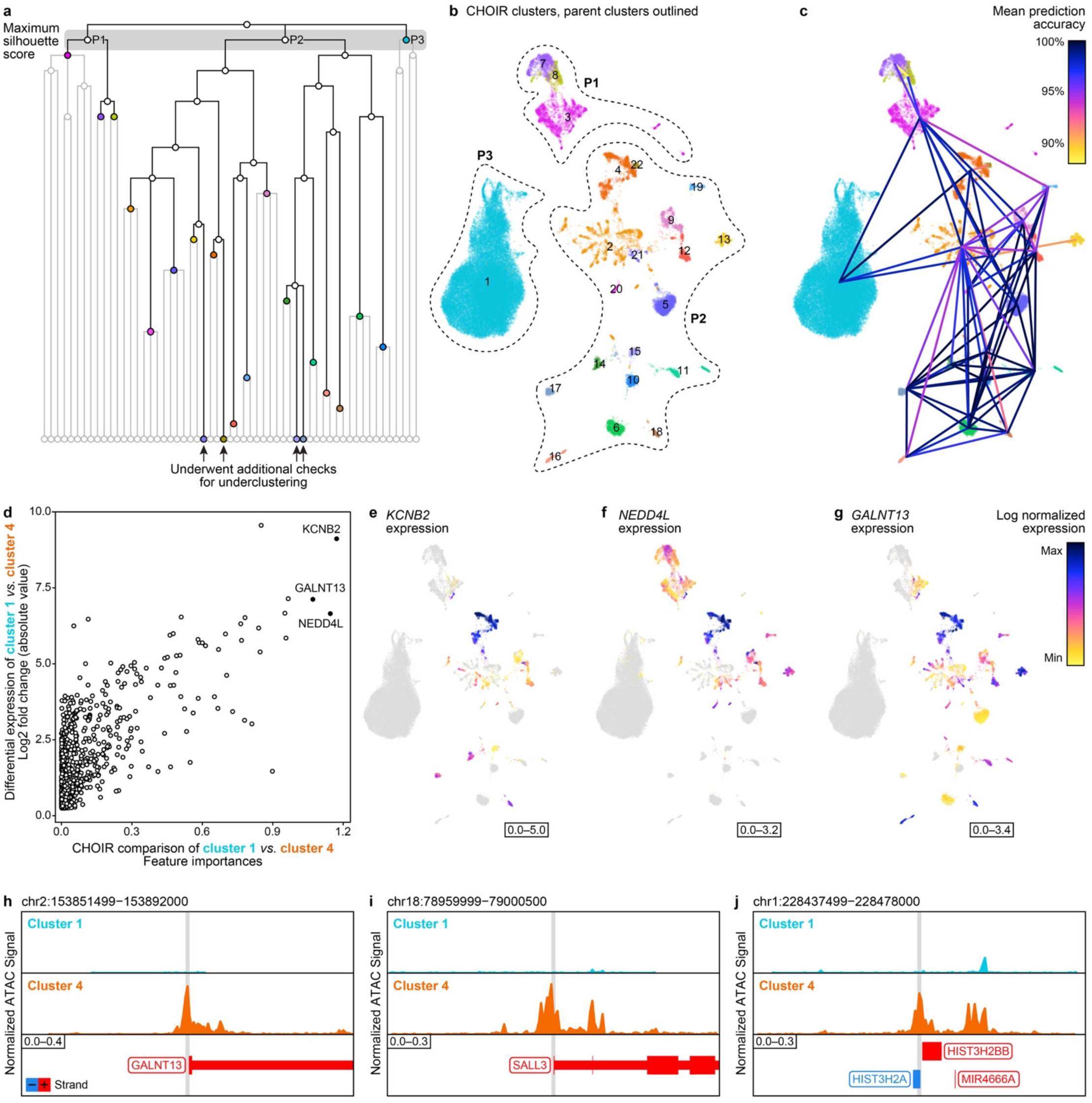
CHOIR effectively clusters multi-omic data and identifies cluster-specific features in both modalities. **a**. Example of a hierarchical clustering tree generated by applying the default parameters of CHOIR to the RNA-seq and ATAC-seq features of the Wang et al. 2022 multi-omic dataset from human retinal cells. Final clusters identified are shown in color, pruned branches are shown in light grey. P1–P3 refer to parent clusters identified by maximizing the silhouette score, highlighted by the grey box. **b.** UMAP embedding colored according to the 22 clusters identified by CHOIR. Parent clusters identified by maximizing the silhouette score are outlined in dashed lines. **c.** Mean prediction accuracy scores indicated by color of lines connecting pairs of final clusters identified by CHOIR. Each line connects the two clusters compared and is colored by the mean prediction accuracy. Not all final cluster pairs were directly compared by permutation testing because some pairs did not meet distance or adjacency criteria. **d.** Dot plot comparing the feature importances extracted from the random forest comparisons computed by CHOIR for cluster 1 (rod cells) versus cluster 4 (cone cells) with the log fold change of gene expression between cluster 1 and cluster 4 identified using Seurat. Each dot represents an individual gene and all genes are shown. **e–g.** UMAP embeddings colored according to the expression level of the three RNA-seq features with the highest feature importances in the comparison of CHOIR cluster 1 versus cluster 4: *KCNB2* (**e**), *NEDD4L* (**f**), and *GALNT13* (**g**) were all enriched in cone cells. **h–j.** Genome track visualizations of the three ATAC-seq features with the highest feature importance in the comparison of CHOIR cluster 1 versus cluster 4: the *GALNT13* locus (chr2:153,851,499–153,892,000) (**h**), *SALL3* locus (chr18:78,959,999–79,000,500) (**i**), and *HIST3H2BB* locus (chr1:228,437,499–228,478,000) (**j**).

**Extended Data Fig. 2:**
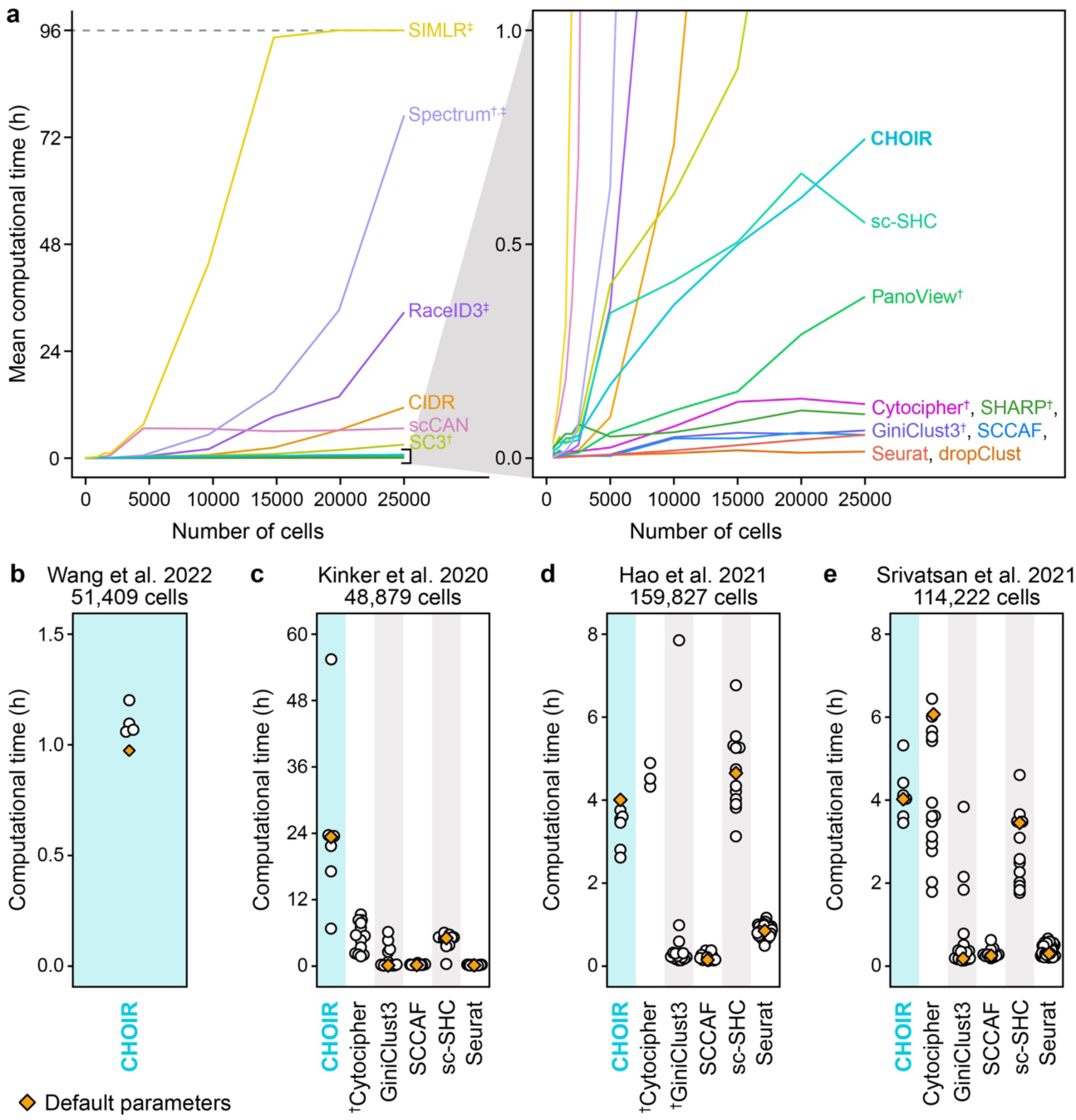
Computational time across clustering methods. **a**. Line plots showing computational time required for the best-performing parameter setting for each of the 15 clustering methods applied to simulated data, averaged across all simulated datasets of each size. Methods with a maximum runtime under one hour are highlighted in the box to the right. Symbols indicate clustering methods that ^†^failed to run or ^‡^did not complete within the maximum allotted runtime of 96 hours for at least one dataset. **b–e.** Computational time of each parameter setting tested for each method for the Wang et al. 2022 multi-omic ATAC-seq and RNA-seq dataset (**b**), Kinker et al. 2020 cancer cell line dataset (**c**), Hao et al. 2021 CITE-seq dataset (**d**), and Srivatsan et al. 2021 sci-Space dataset (**e**). Symbols indicate clustering methods in which one or more parameter settings ^†^failed to run.

**Extended Data Fig. 3:**
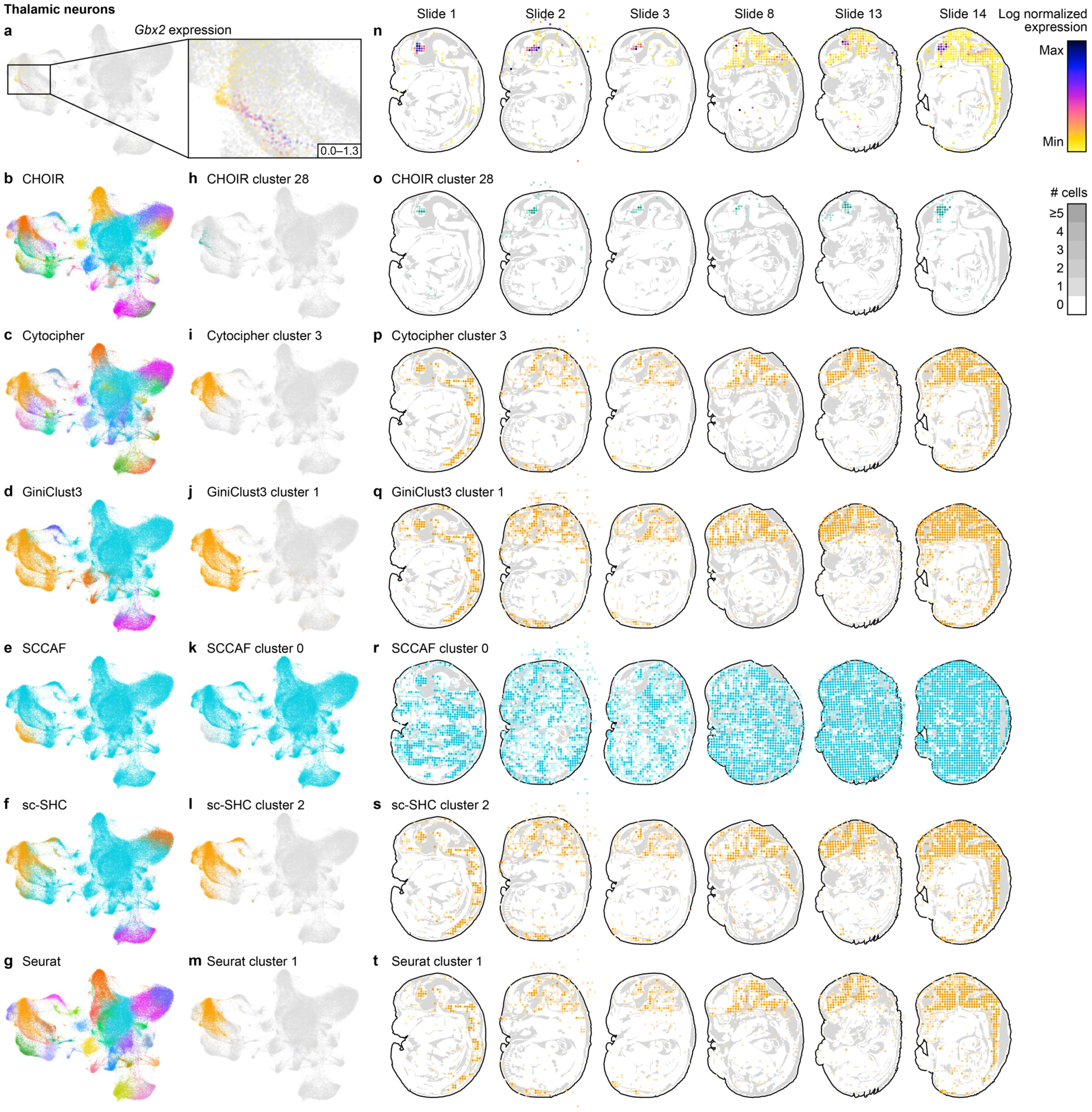
CHOIR cluster 28 represents a population of neurons localized within the developing thalamus characterized by high *Gbx2* expression. **a–g**. UMAP embedding of the Srivatsan et al. 2021 whole mouse embryo sci-Space dataset, colored according to the expression level of thalamic marker gene *Gbx2* (**a**), or the clusters identified by the default parameters of CHOIR (**b**), Cytocipher (**c**), GiniClust3 (**d**), SCCAF (**e**), sc-SHC (**f**), or Seurat (**g**). A zoom in of *Gbx2* expression is shown to the right of panel (**a**). **h–m.** UMAP embedding of the Srivatsan et al. 2021 whole mouse embryo sci-Space dataset, colored by the cluster shown in panels (**a–g**) that harbors the *Gbx2*-expressing thalamic neurons for each of the methods shown to the left. **n.** Distribution of thalamic marker gene *Gbx2* in all sections with >25 cells belonging to CHOIR cluster 28 (thalamic neurons). **o–t.** Distribution of the method-specific cluster shown to the left in panels (**h–m**) across all sections shown in panel (**n**). The darker the shade of the cluster color, the more cells of the respective cluster were detected at the indicated location.

## SUPPLEMENTARY INFORMATION FOR

**Supplementary Fig. 1:**
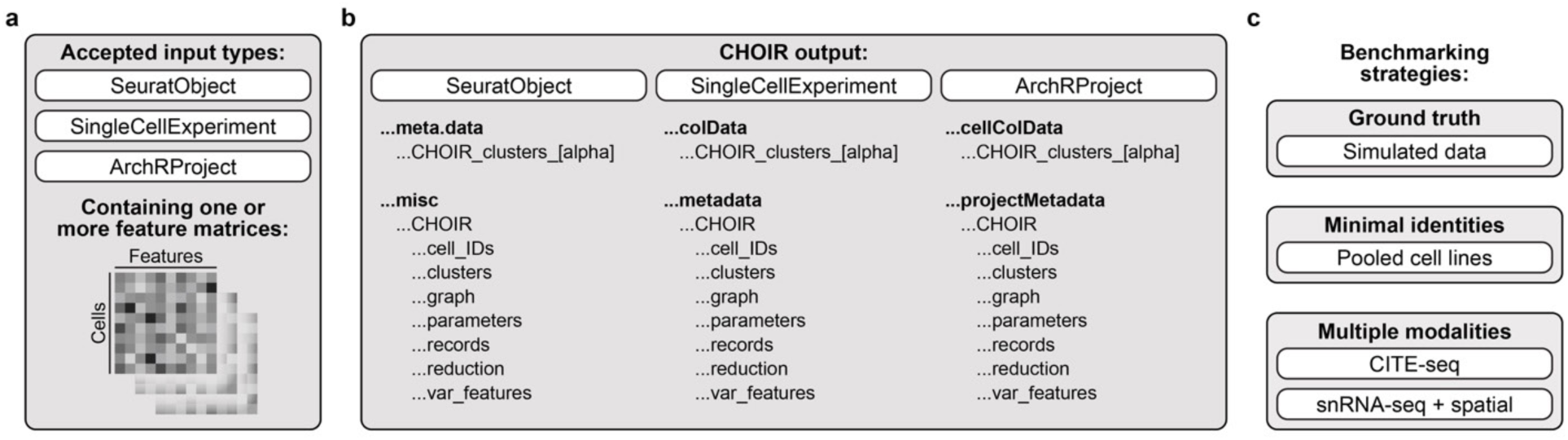
CHOIR provides compatibility for multiple data types, formats, and analysis outputs. **a**. Schematic showing the types of input data accepted by CHOIR. **b.** Schematic showing the CHOIR output locations by object type. **c.** A three-tiered benchmarking approach was used to assess the performance of CHOIR relative to existing clustering methods.

**Supplementary Fig. 2:**
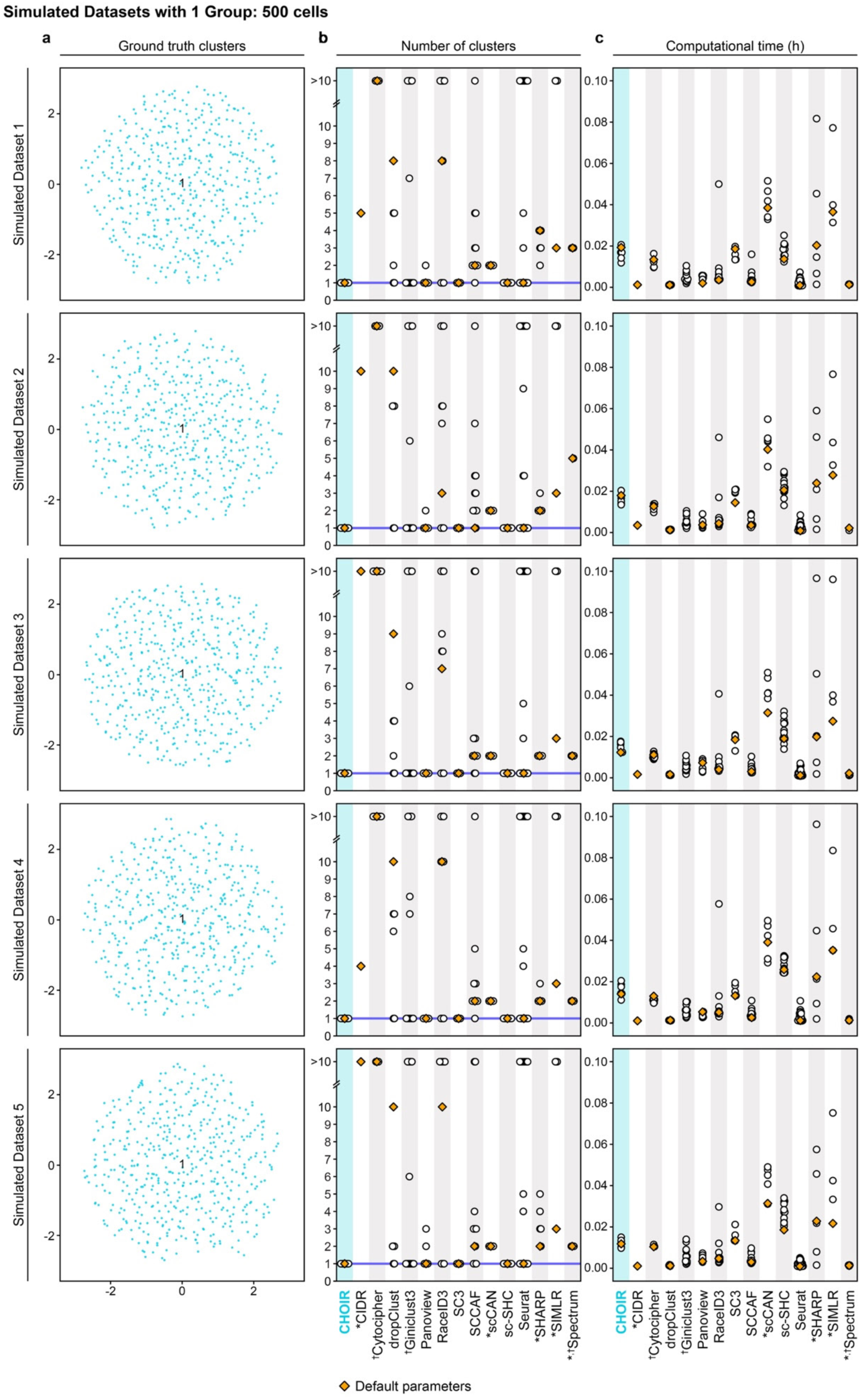
Detailed benchmarking results from simulated datasets with one ground truth group. **a**. UMAP embeddings of each dataset colored according to the ground truth groups. **b.** Number of clusters of each parameter setting tested for each method. Symbols indicate clustering methods that *assumed ≥2 clusters, had one or more parameter settings ^†^failed to run, or ^‡^had one or more parameter settings that did not complete within the maximum allotted runtime of 96 hours for at least one dataset. **c.** Computational time of each parameter setting tested for each method.

**Supplementary Fig. 3:**
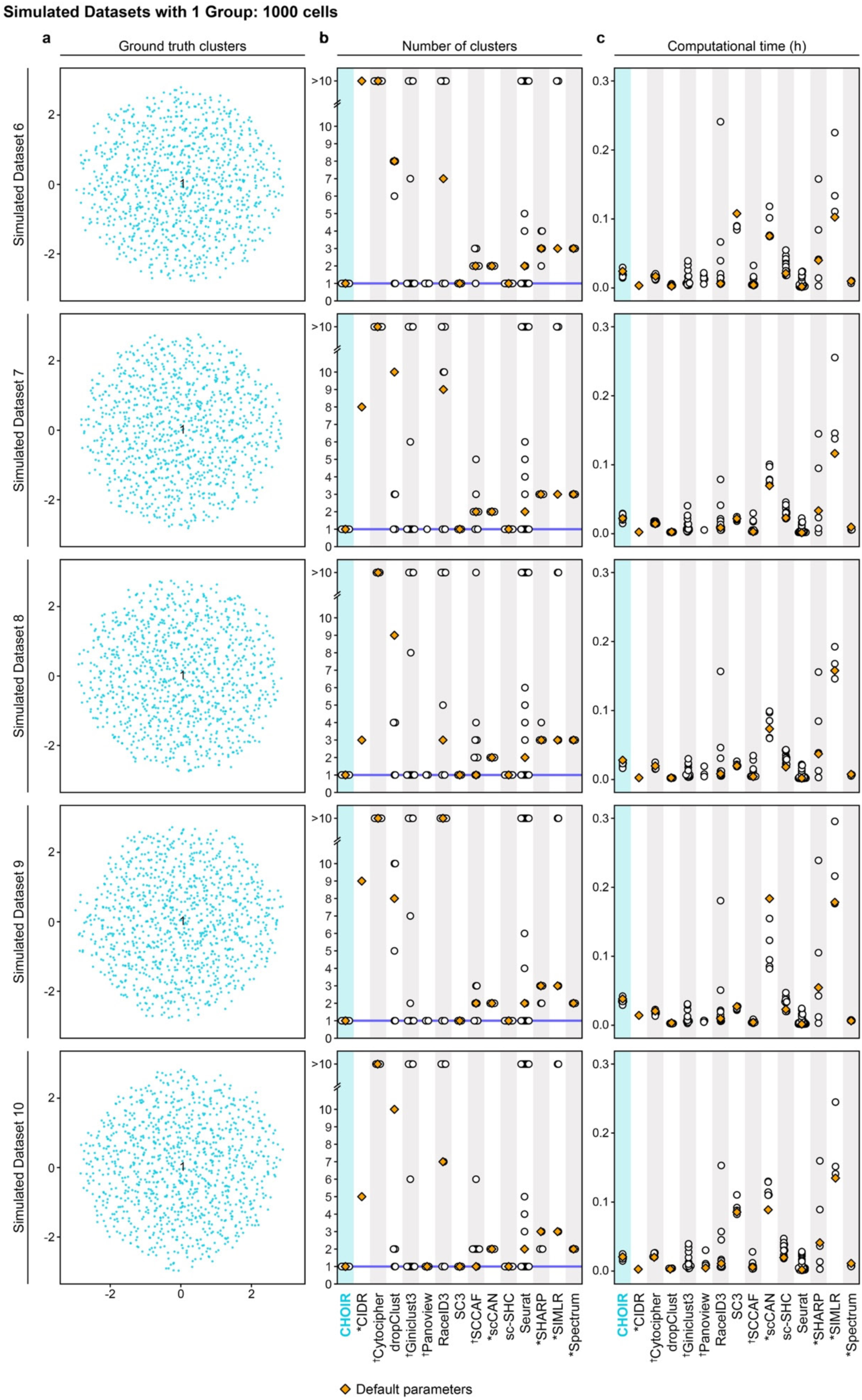
Detailed benchmarking results from simulated datasets with one ground truth group. **a**. UMAP embeddings of each dataset colored according to the ground truth groups. **b.** Number of clusters of each parameter setting tested for each method. Symbols indicate clustering methods that *assumed ≥2 clusters, had one or more parameter settings ^†^failed to run, or ^‡^had one or more parameter settings that did not complete within the maximum allotted runtime of 96 hours for at least one dataset. **c.** Computational time of each parameter setting tested for each method.

**Supplementary Fig. 4:**
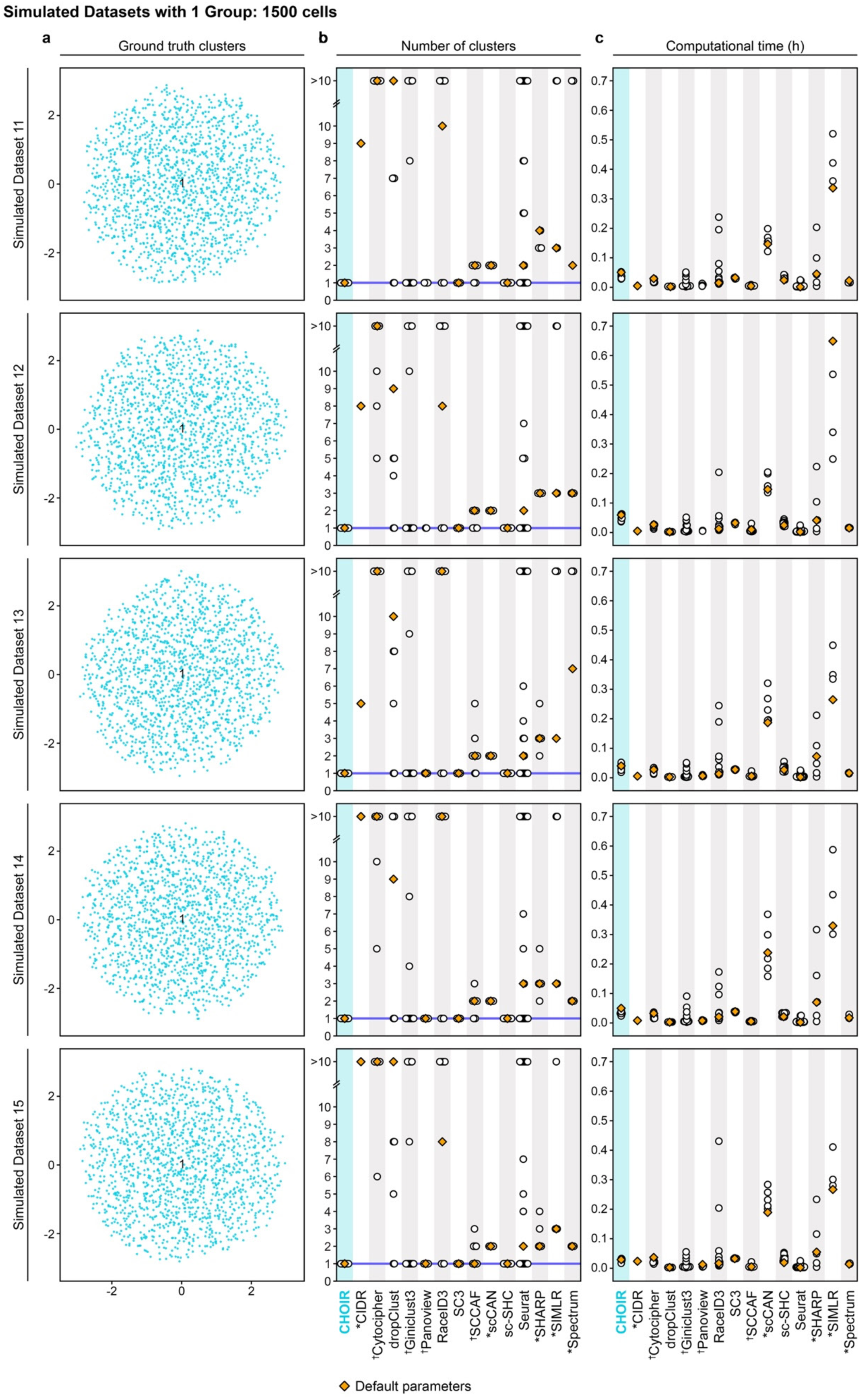
Detailed benchmarking results from simulated datasets with one ground truth group. **a**. UMAP embeddings of each dataset colored according to the ground truth groups. **b.** Number of clusters of each parameter setting tested for each method. Symbols indicate clustering methods that *assumed ≥2 clusters, had one or more parameter settings ^†^failed to run, or ^‡^had one or more parameter settings that did not complete within the maximum allotted runtime of 96 hours for at least one dataset. **c.** Computational time of each parameter setting tested for each method.

**Supplementary Fig. 5:**
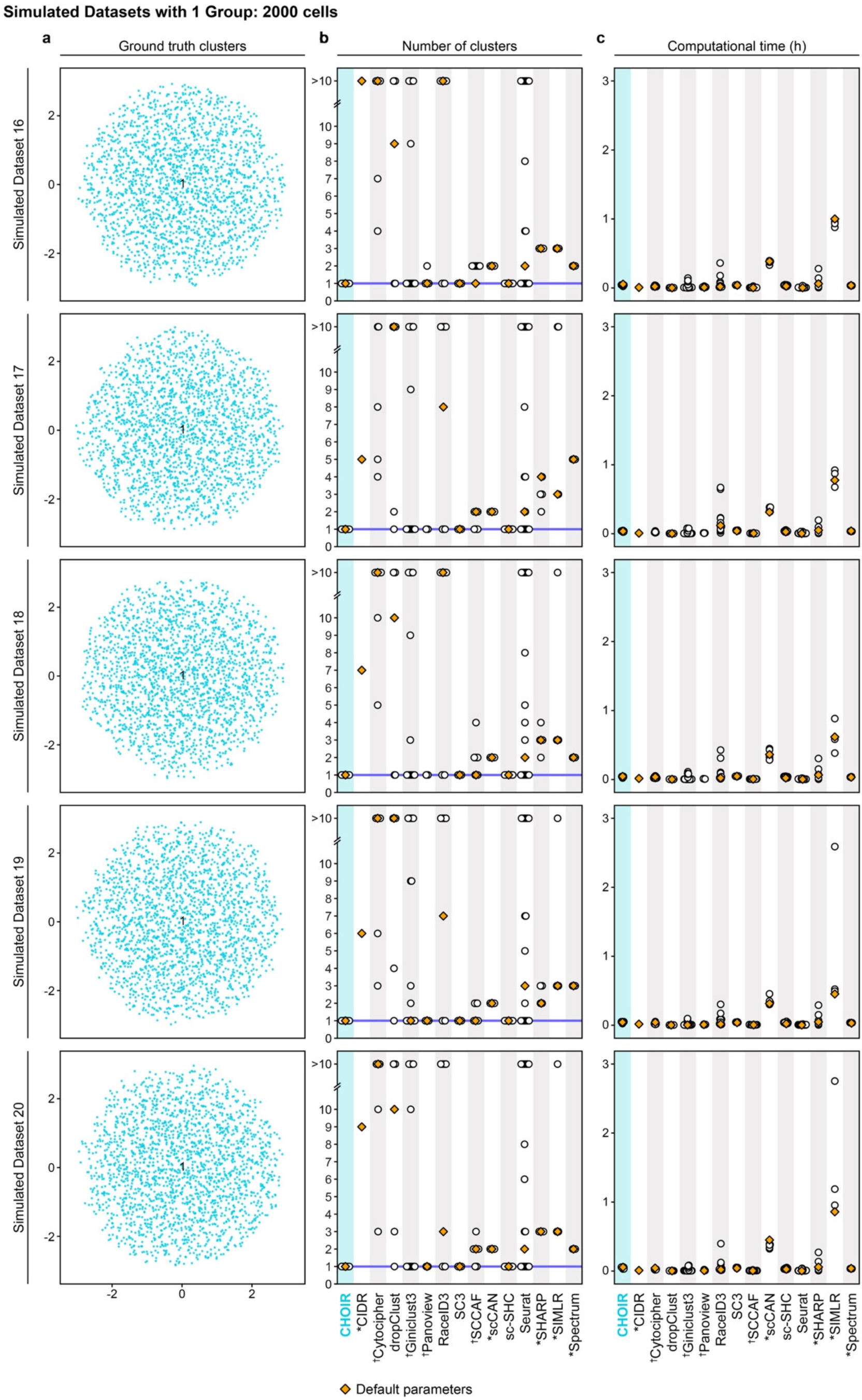
Detailed benchmarking results from simulated datasets with one ground truth group. **a**. UMAP embeddings of each dataset colored according to the ground truth groups. **b.** Number of clusters of each parameter setting tested for each method. Symbols indicate clustering methods that *assumed ≥2 clusters, had one or more parameter settings ^†^failed to run, or ^‡^had one or more parameter settings that did not complete within the maximum allotted runtime of 96 hours for at least one dataset. **c.** Computational time of each parameter setting tested for each method.

**Supplementary Fig. 6:**
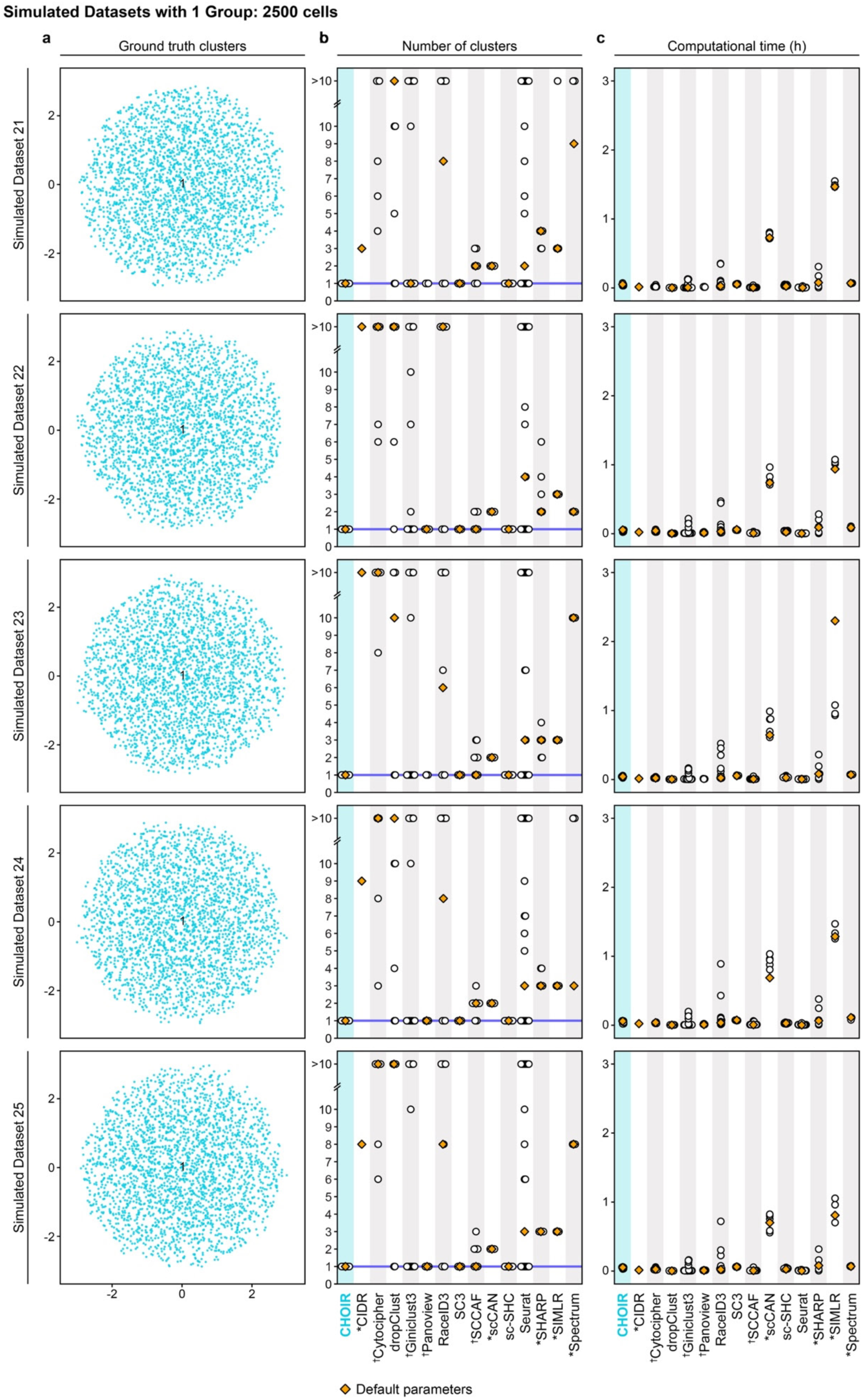
Detailed benchmarking results from simulated datasets with one ground truth group. **a**. UMAP embeddings of each dataset colored according to the ground truth groups. **b.** Number of clusters of each parameter setting tested for each method. Symbols indicate clustering methods that *assumed ≥2 clusters, had one or more parameter settings ^†^failed to run, or ^‡^had one or more parameter settings that did not complete within the maximum allotted runtime of 96 hours for at least one dataset. **c.** Computational time of each parameter setting tested for each method.

**Supplementary Fig. 7:**
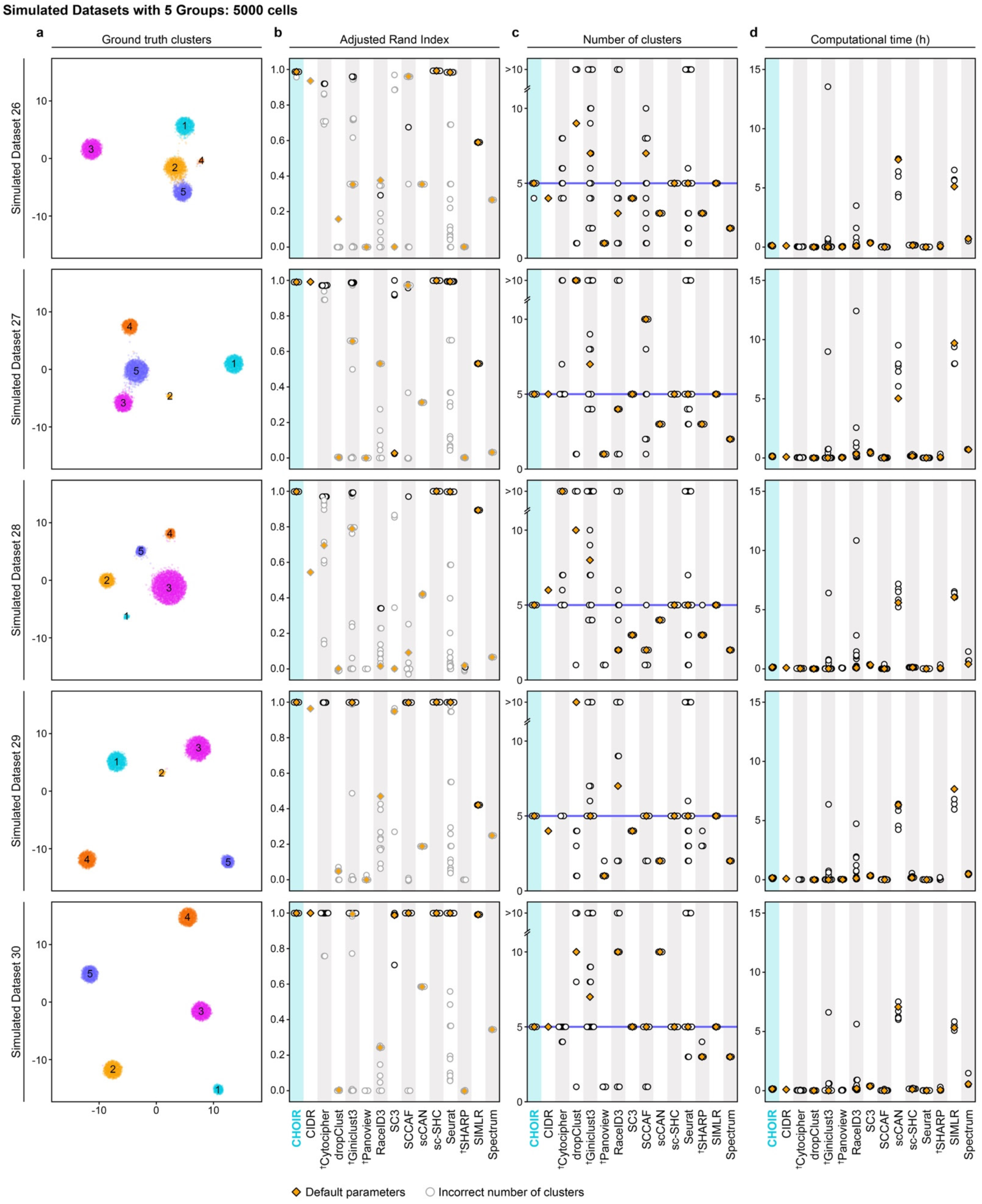
Detailed benchmarking results from simulated datasets with 5–20 ground truth groups. **a**. UMAP embeddings of each dataset colored according to the ground truth groups. **b.** Adjusted Rand Index of each parameter setting tested for each method. Symbols indicate clustering methods for which one or more parameter settings ^†^failed to run or ^‡^did not complete within the maximum allotted runtime of 96 hours for at least one dataset. **c.** Number of clusters of each parameter setting tested for each method. **d.** Computational time of each parameter setting tested for each method.

**Supplementary Fig. 8:**
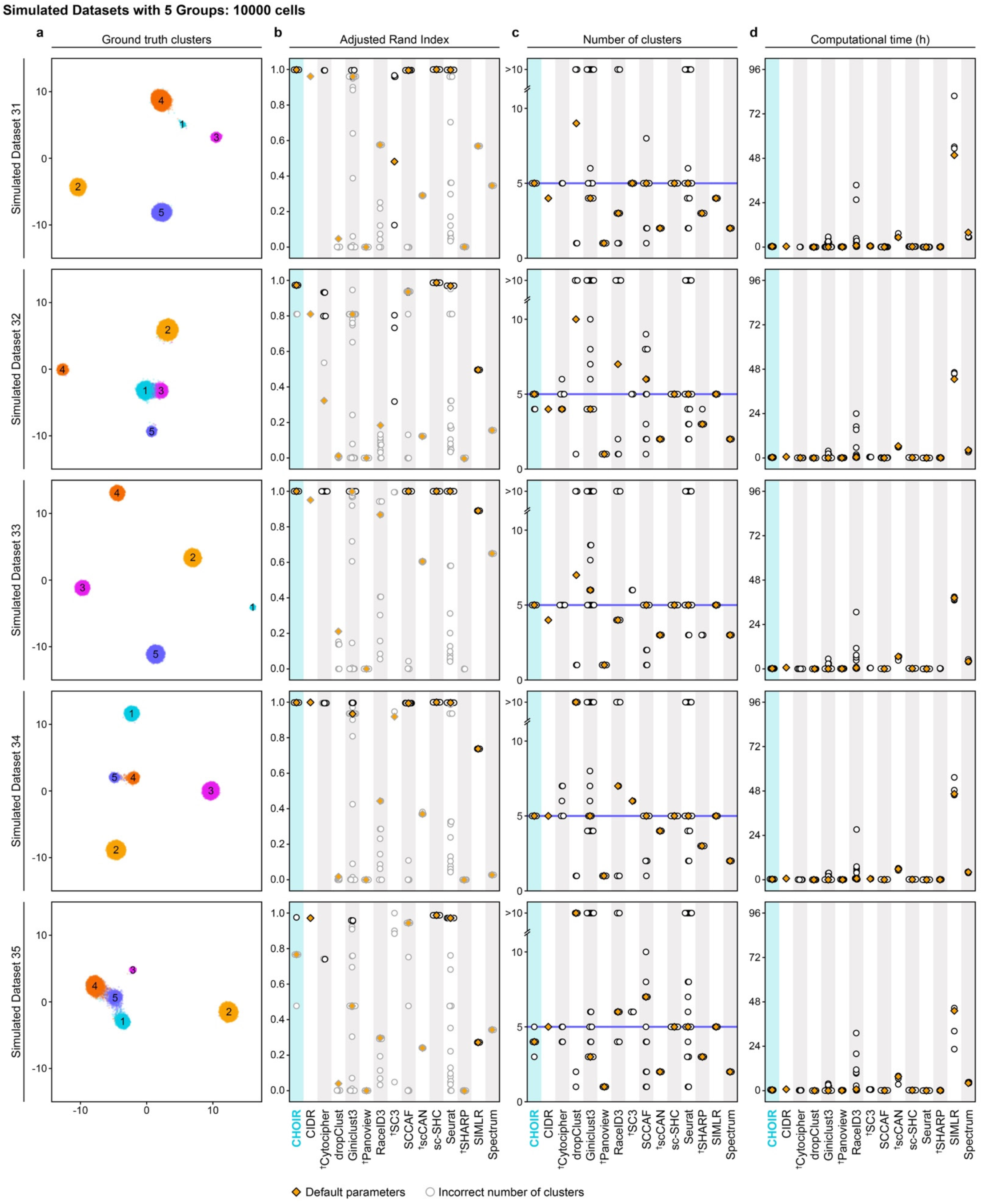
Detailed benchmarking results from simulated datasets with 5–20 ground truth groups. **a**. UMAP embeddings of each dataset colored according to the ground truth groups. **b.** Adjusted Rand Index of each parameter setting tested for each method. Symbols indicate clustering methods for which one or more parameter settings ^†^failed to run or ^‡^did not complete within the maximum allotted runtime of 96 hours for at least one dataset. **c.** Number of clusters of each parameter setting tested for each method. **d.** Computational time of each parameter setting tested for each method.

**Supplementary Fig. 9:**
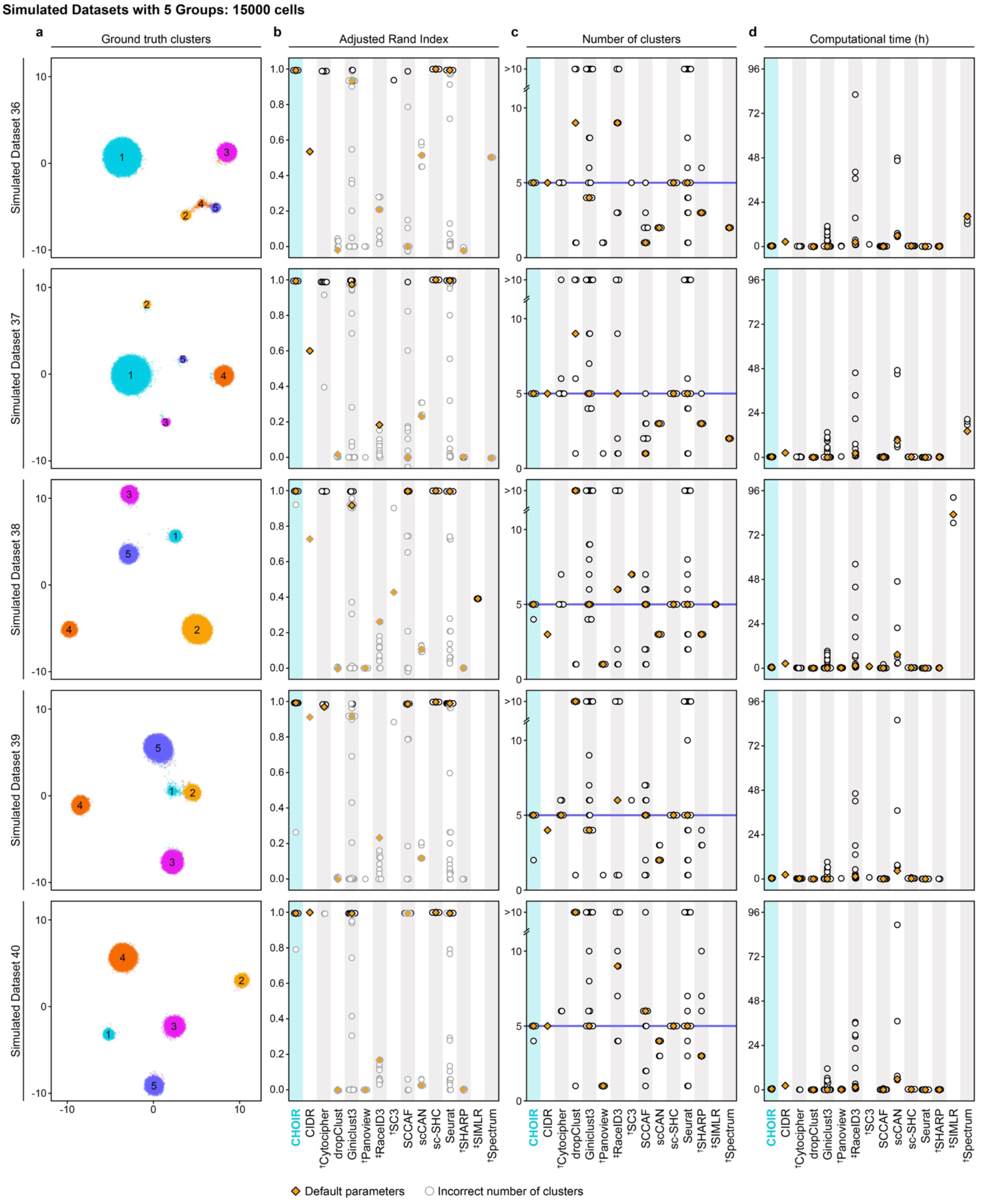
Detailed benchmarking results from simulated datasets with 5–20 ground truth groups. **a**. UMAP embeddings of each dataset colored according to the ground truth groups. **b.** Adjusted Rand Index of each parameter setting tested for each method. Symbols indicate clustering methods for which one or more parameter settings ^†^failed to run or ^‡^did not complete within the maximum allotted runtime of 96 hours for at least one dataset. **c.** Number of clusters of each parameter setting tested for each method. **d.** Computational time of each parameter setting tested for each method.

**Supplementary Fig. 10:**
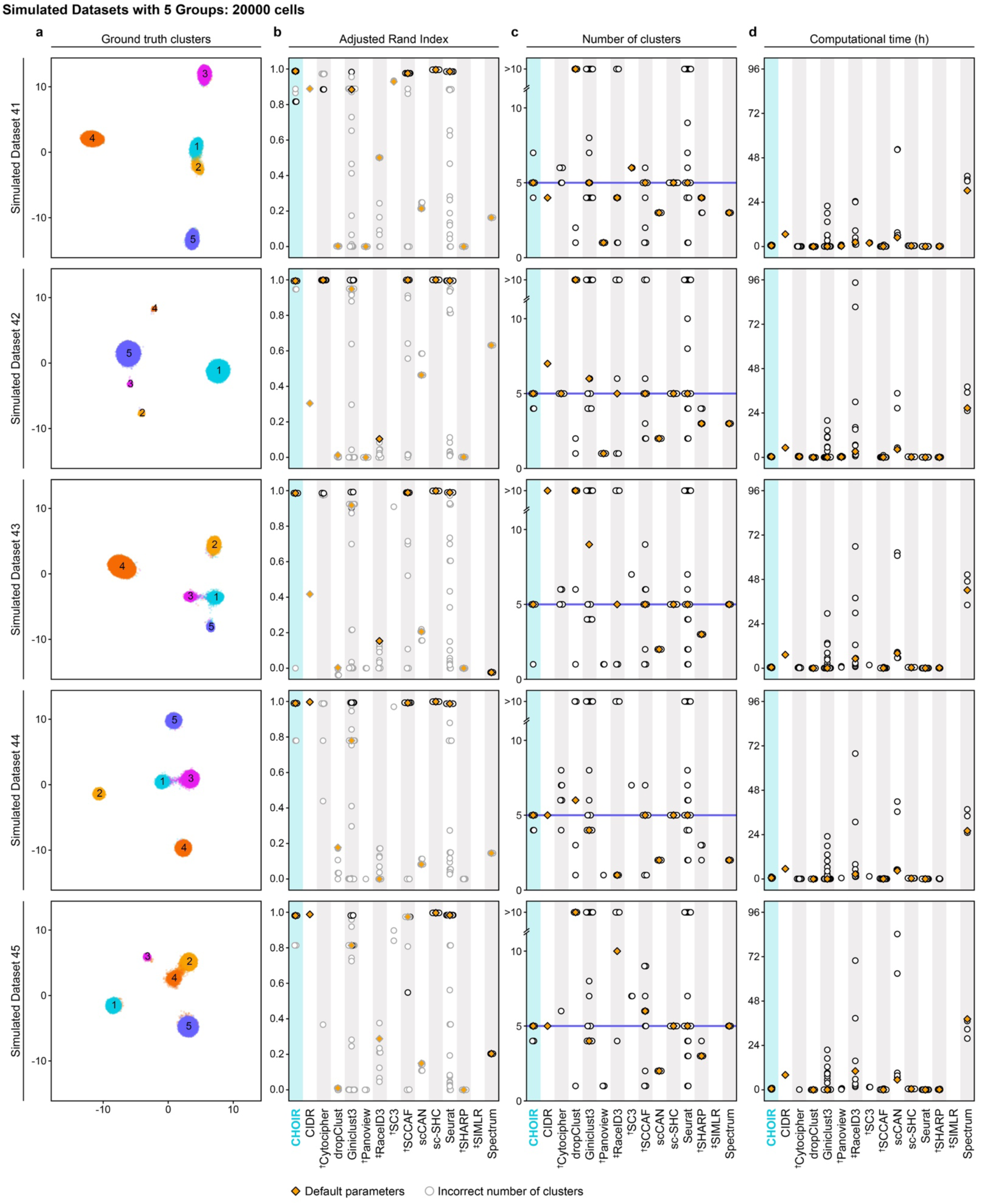
Detailed benchmarking results from simulated datasets with 5–20 ground truth groups. **a**. UMAP embeddings of each dataset colored according to the ground truth groups. **b.** Adjusted Rand Index of each parameter setting tested for each method. Symbols indicate clustering methods for which one or more parameter settings ^†^failed to run or ^‡^did not complete within the maximum allotted runtime of 96 hours for at least one dataset. **c.** Number of clusters of each parameter setting tested for each method. **d.** Computational time of each parameter setting tested for each method.

**Supplementary Fig. 11:**
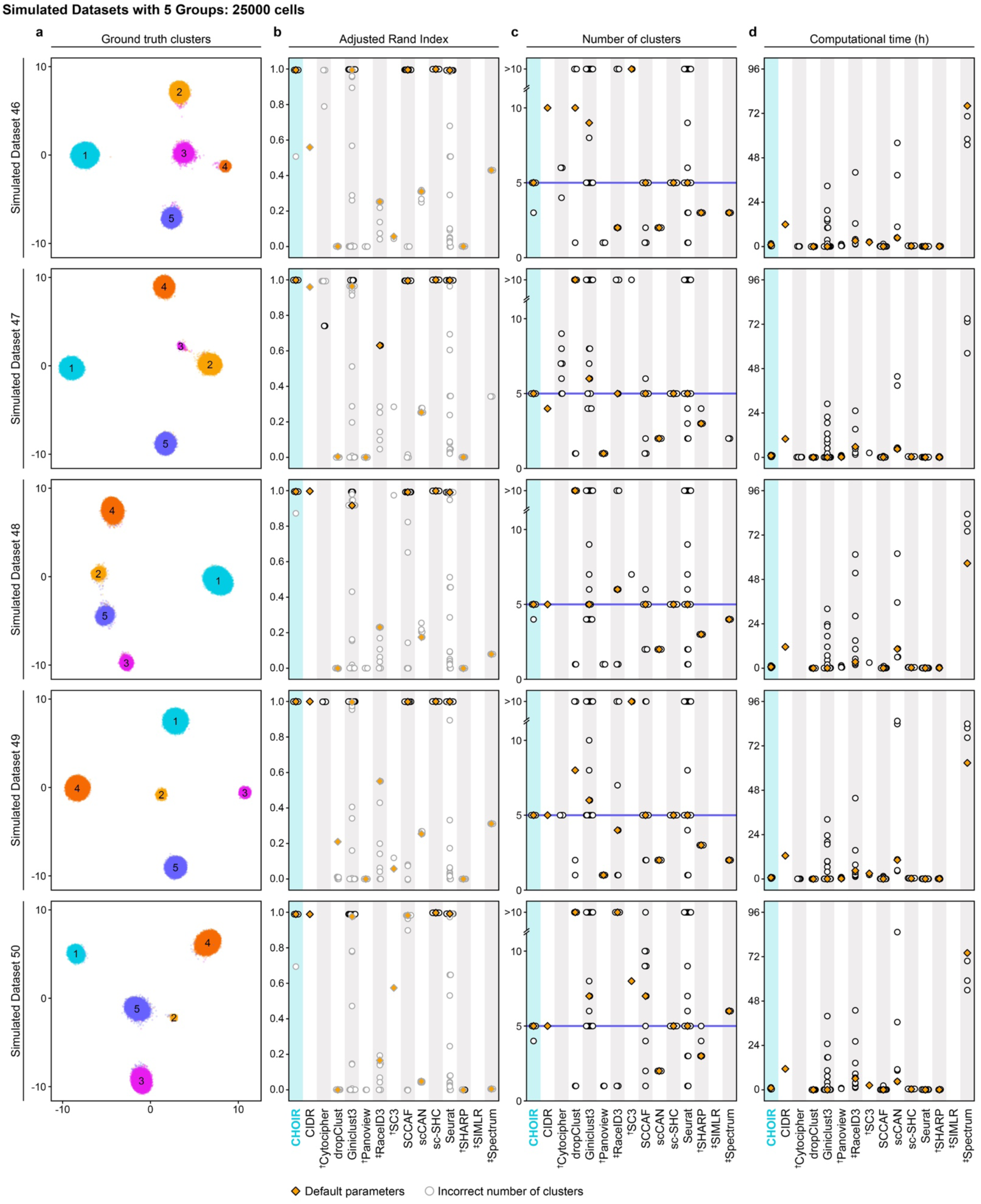
Detailed benchmarking results from simulated datasets with 5–20 ground truth groups. **a**. UMAP embeddings of each dataset colored according to the ground truth groups. **b.** Adjusted Rand Index of each parameter setting tested for each method. Symbols indicate clustering methods for which one or more parameter settings ^†^failed to run or ^‡^did not complete within the maximum allotted runtime of 96 hours for at least one dataset. **c.** Number of clusters of each parameter setting tested for each method. **d.** Computational time of each parameter setting tested for each method.

**Supplementary Fig. 12:**
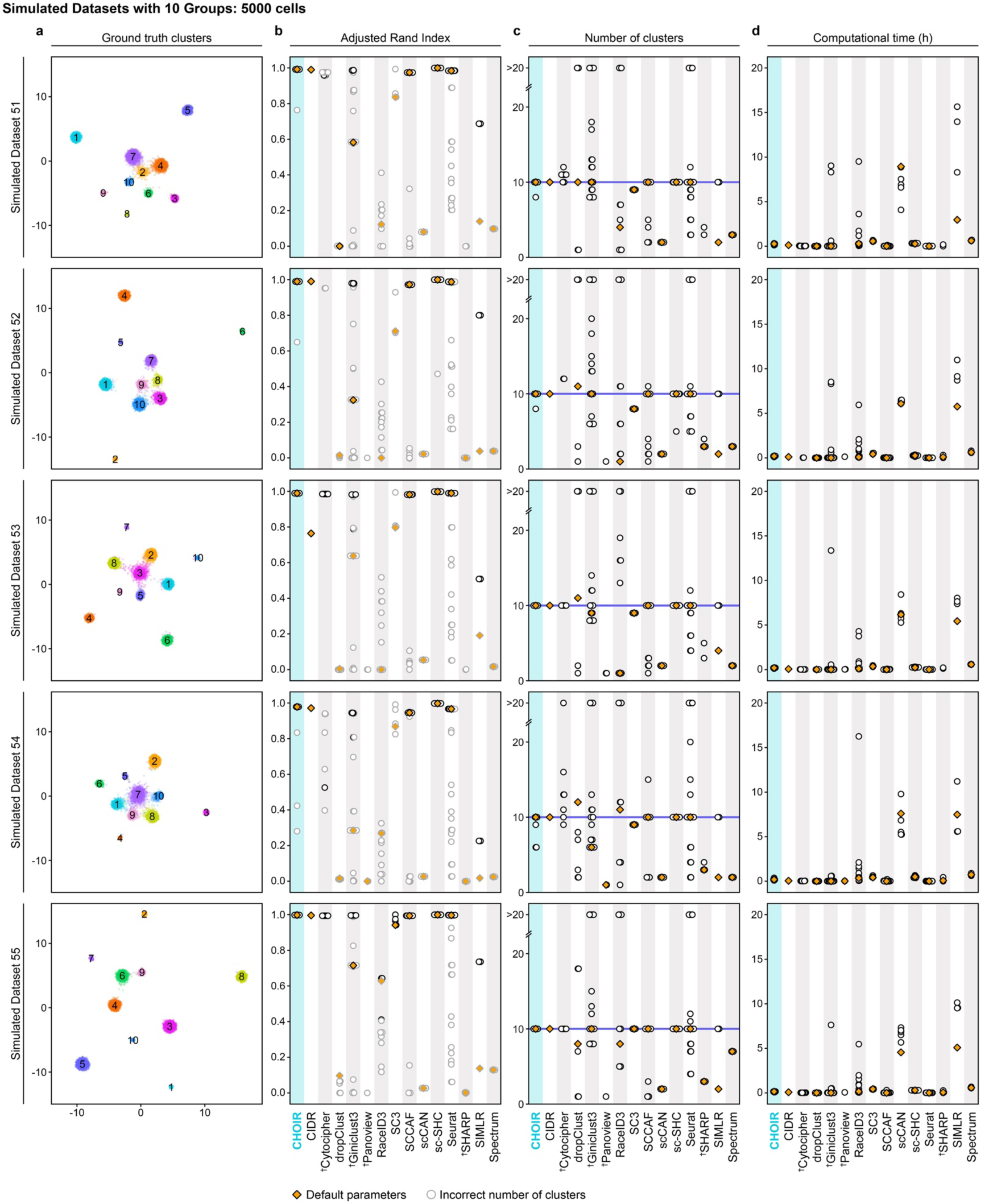
Detailed benchmarking results from simulated datasets with 5–20 ground truth groups. **a**. UMAP embeddings of each dataset colored according to the ground truth groups. **b.** Adjusted Rand Index of each parameter setting tested for each method. Symbols indicate clustering methods for which one or more parameter settings ^†^failed to run or ^‡^did not complete within the maximum allotted runtime of 96 hours for at least one dataset. **c.** Number of clusters of each parameter setting tested for each method. **d.** Computational time of each parameter setting tested for each method.

**Supplementary Fig. 13:**
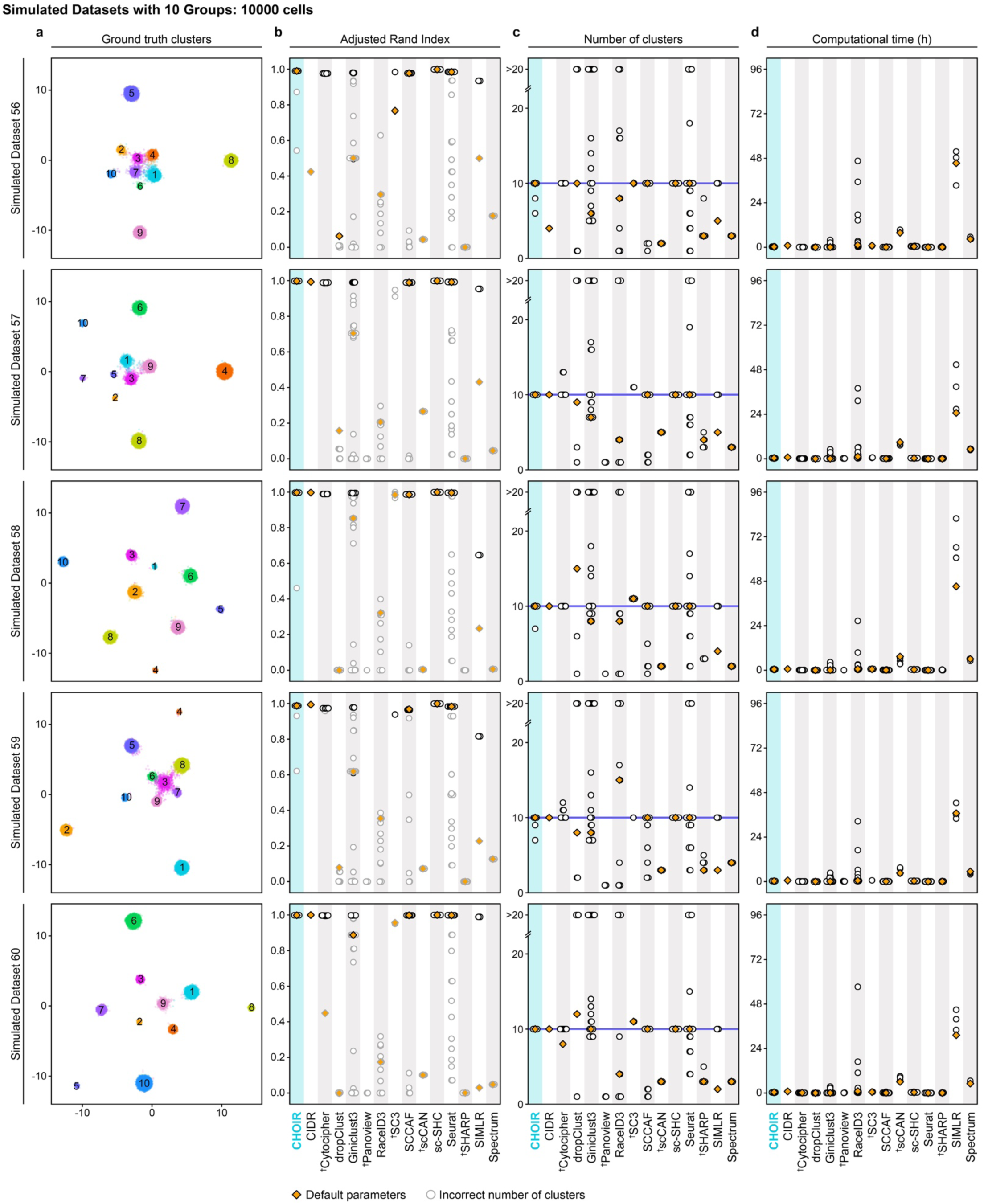
Detailed benchmarking results from simulated datasets with 5–20 ground truth groups. **a**. UMAP embeddings of each dataset colored according to the ground truth groups. **b.** Adjusted Rand Index of each parameter setting tested for each method. Symbols indicate clustering methods for which one or more parameter settings ^†^failed to run or ^‡^did not complete within the maximum allotted runtime of 96 hours for at least one dataset. **c.** Number of clusters of each parameter setting tested for each method. **d.** Computational time of each parameter setting tested for each method.

**Supplementary Fig. 14:**
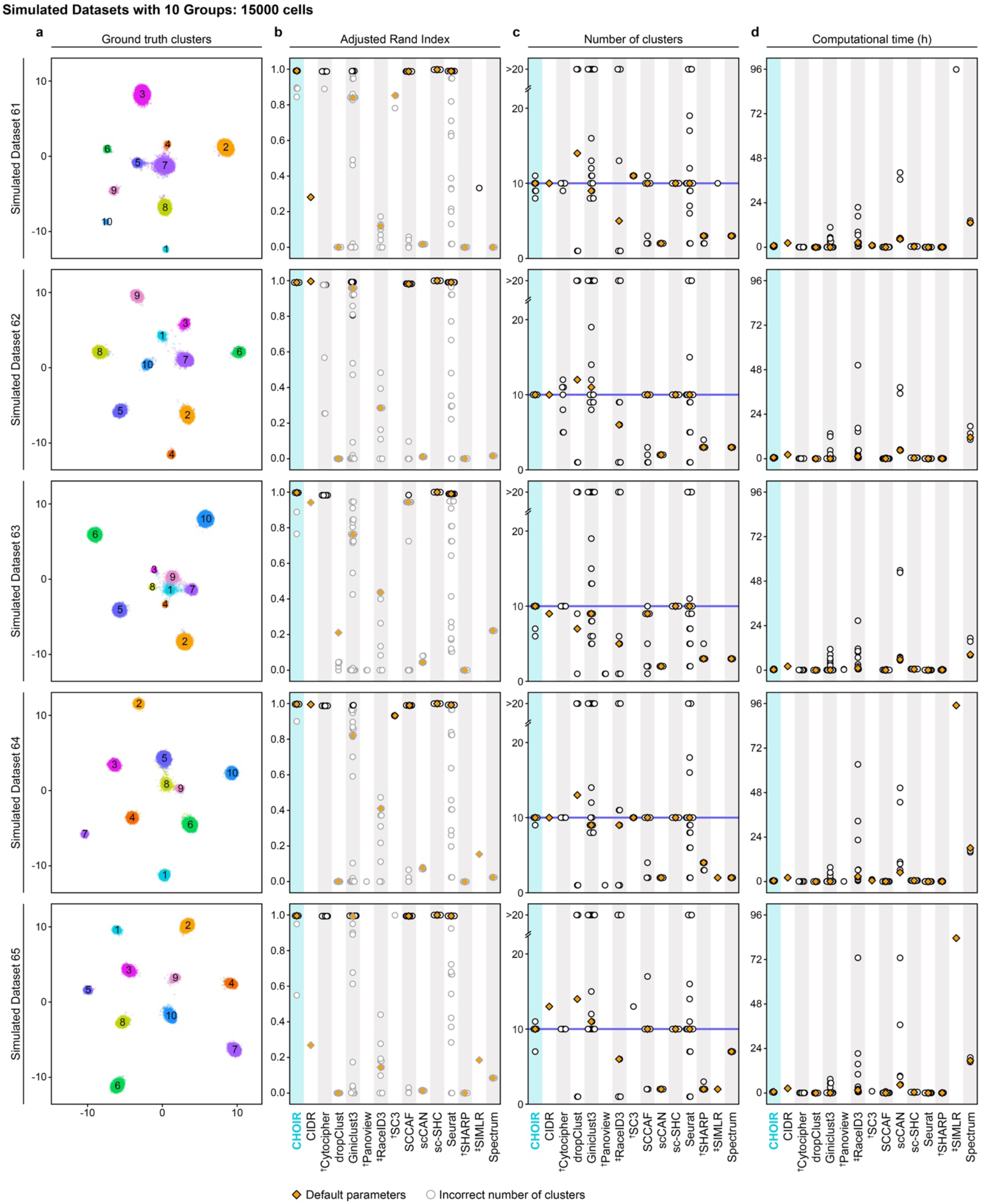
Detailed benchmarking results from simulated datasets with 5–20 ground truth groups. **a**. UMAP embeddings of each dataset colored according to the ground truth groups. **b.** Adjusted Rand Index of each parameter setting tested for each method. Symbols indicate clustering methods for which one or more parameter settings ^†^failed to run or ^‡^did not complete within the maximum allotted runtime of 96 hours for at least one dataset. **c.** Number of clusters of each parameter setting tested for each method. **d.** Computational time of each parameter setting tested for each method.

**Supplementary Fig. 15:**
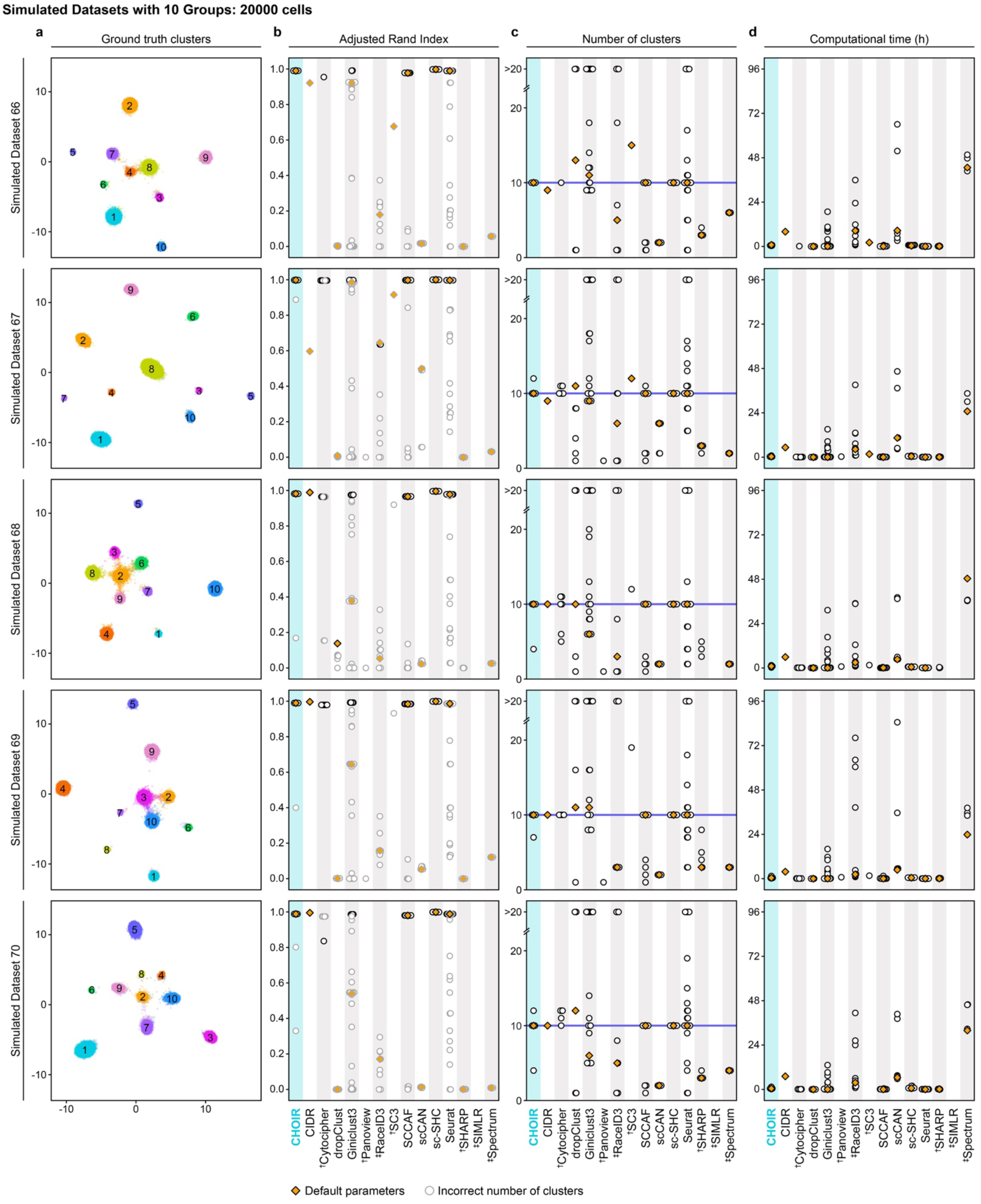
Detailed benchmarking results from simulated datasets with 5–20 ground truth groups. **a**. UMAP embeddings of each dataset colored according to the ground truth groups. **b.** Adjusted Rand Index of each parameter setting tested for each method. Symbols indicate clustering methods for which one or more parameter settings ^†^failed to run or ^‡^did not complete within the maximum allotted runtime of 96 hours for at least one dataset. **c.** Number of clusters of each parameter setting tested for each method. **d.** Computational time of each parameter setting tested for each method.

**Supplementary Fig. 16:**
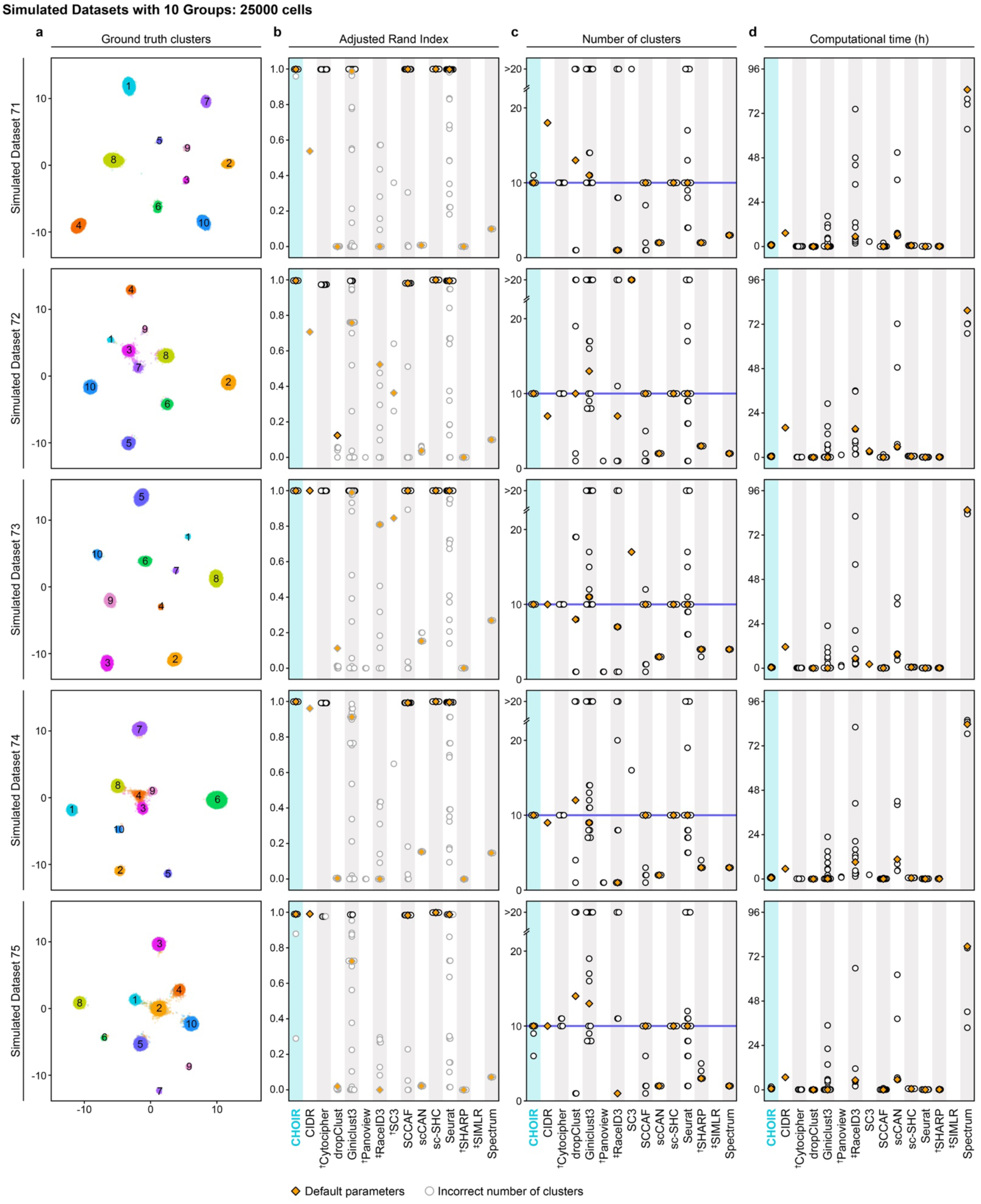
Detailed benchmarking results from simulated datasets with 5–20 ground truth groups. **a**. UMAP embeddings of each dataset colored according to the ground truth groups. **b.** Adjusted Rand Index of each parameter setting tested for each method. Symbols indicate clustering methods for which one or more parameter settings ^†^failed to run or ^‡^did not complete within the maximum allotted runtime of 96 hours for at least one dataset. **c.** Number of clusters of each parameter setting tested for each method. **d.** Computational time of each parameter setting tested for each method.

**Supplementary Fig. 17:**
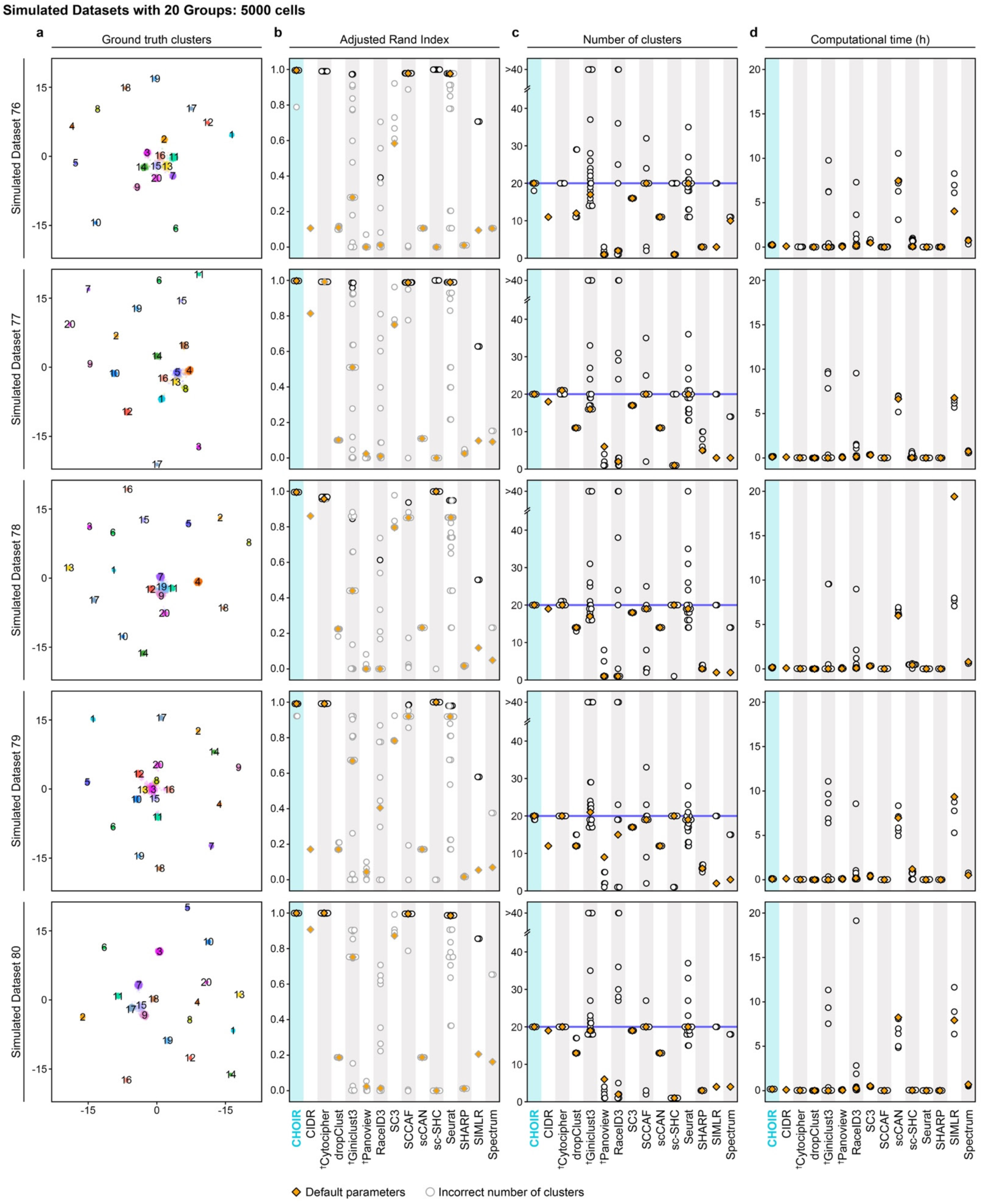
Detailed benchmarking results from simulated datasets with 5–20 ground truth groups. **a**. UMAP embeddings of each dataset colored according to the ground truth groups. **b.** Adjusted Rand Index of each parameter setting tested for each method. Symbols indicate clustering methods for which one or more parameter settings ^†^failed to run or ^‡^did not complete within the maximum allotted runtime of 96 hours for at least one dataset. **c.** Number of clusters of each parameter setting tested for each method. **d.** Computational time of each parameter setting tested for each method.

**Supplementary Fig. 18:**
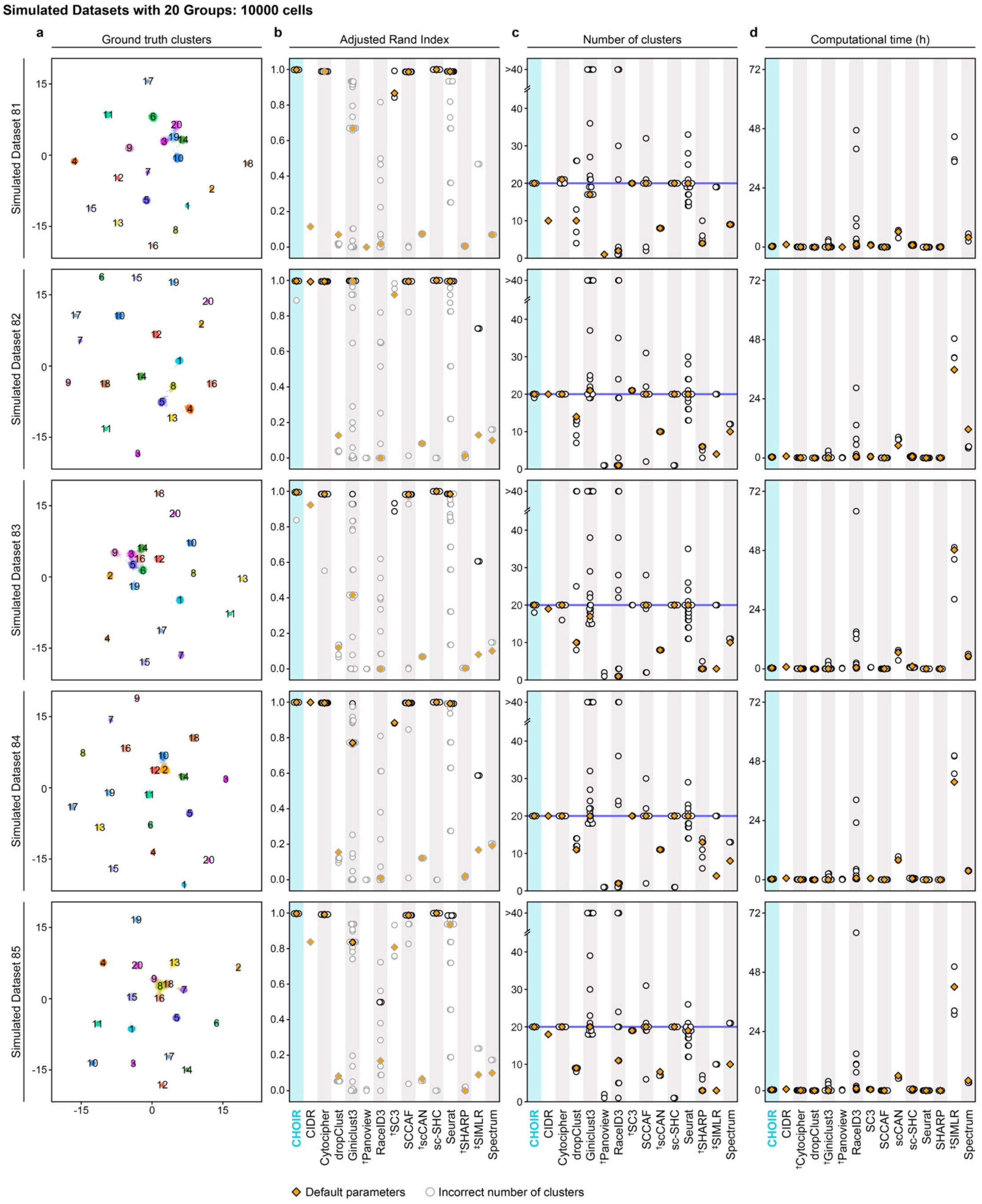
Detailed benchmarking results from simulated datasets with 5–20 ground truth groups. **a**. UMAP embeddings of each dataset colored according to the ground truth groups. **b.** Adjusted Rand Index of each parameter setting tested for each method. Symbols indicate clustering methods for which one or more parameter settings ^†^failed to run or ^‡^did not complete within the maximum allotted runtime of 96 hours for at least one dataset. **c.** Number of clusters of each parameter setting tested for each method. **d.** Computational time of each parameter setting tested for each method.

**Supplementary Fig. 19:**
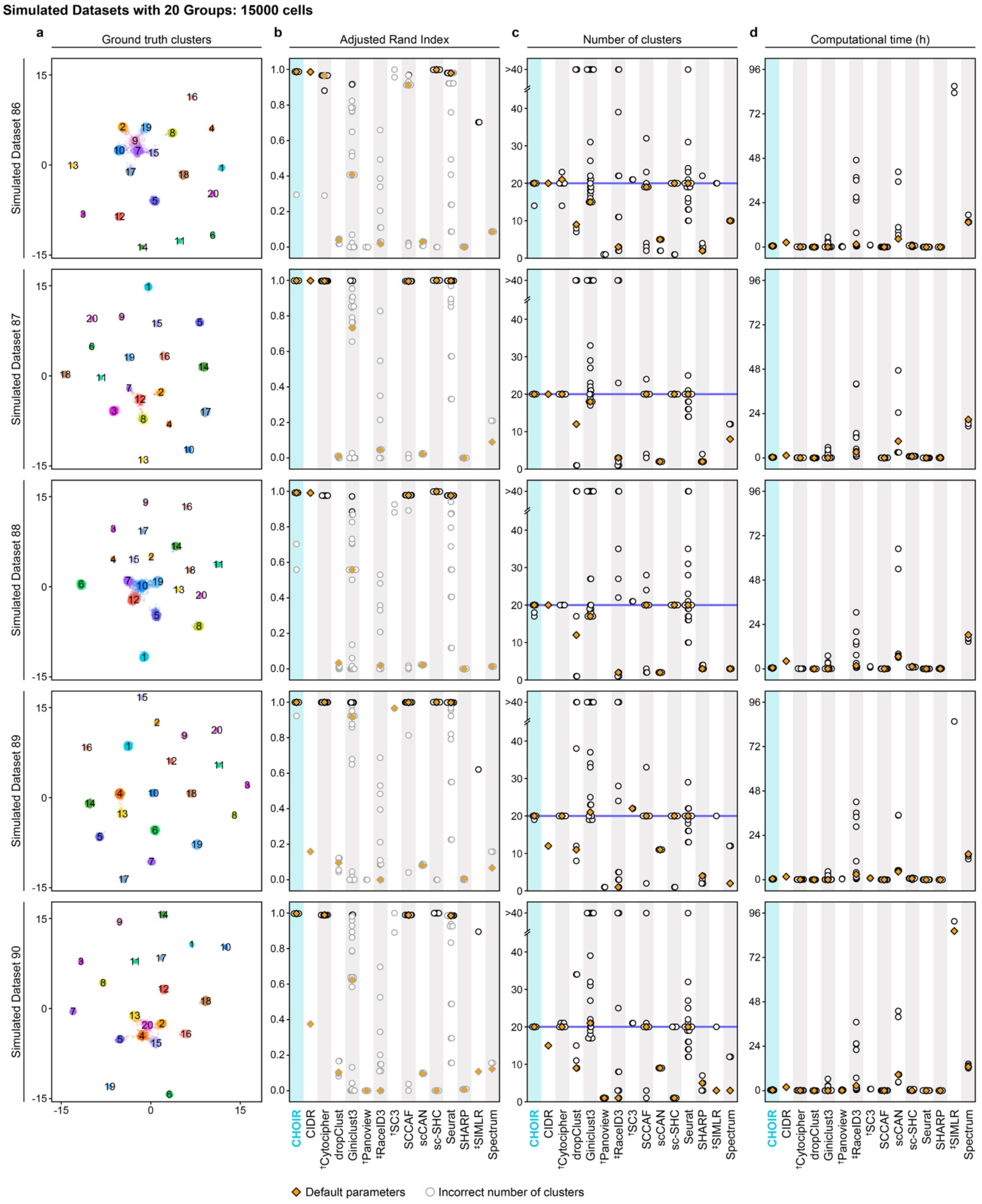
Detailed benchmarking results from simulated datasets with 5–20 ground truth groups. **a**. UMAP embeddings of each dataset colored according to the ground truth groups. **b.** Adjusted Rand Index of each parameter setting tested for each method. Symbols indicate clustering methods for which one or more parameter settings ^†^failed to run or ^‡^did not complete within the maximum allotted runtime of 96 hours for at least one dataset. **c.** Number of clusters of each parameter setting tested for each method. **d.** Computational time of each parameter setting tested for each method.

**Supplementary Fig. 20:**
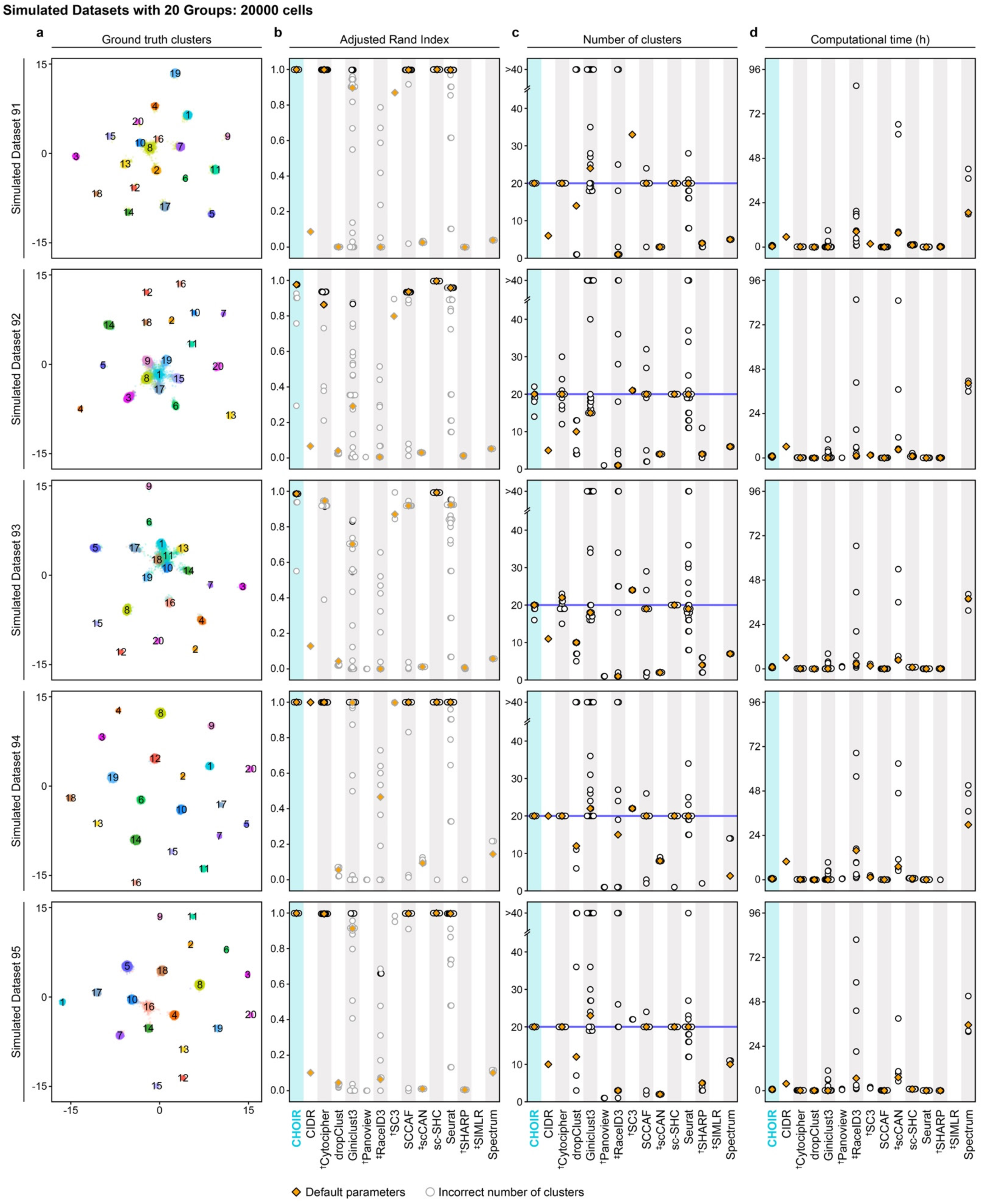
Detailed benchmarking results from simulated datasets with 5–20 ground truth groups. **a**. UMAP embeddings of each dataset colored according to the ground truth groups. **b.** Adjusted Rand Index of each parameter setting tested for each method. Symbols indicate clustering methods for which one or more parameter settings ^†^failed to run or ^‡^did not complete within the maximum allotted runtime of 96 hours for at least one dataset. **c.** Number of clusters of each parameter setting tested for each method. **d.** Computational time of each parameter setting tested for each method.

**Supplementary Fig. 21:**
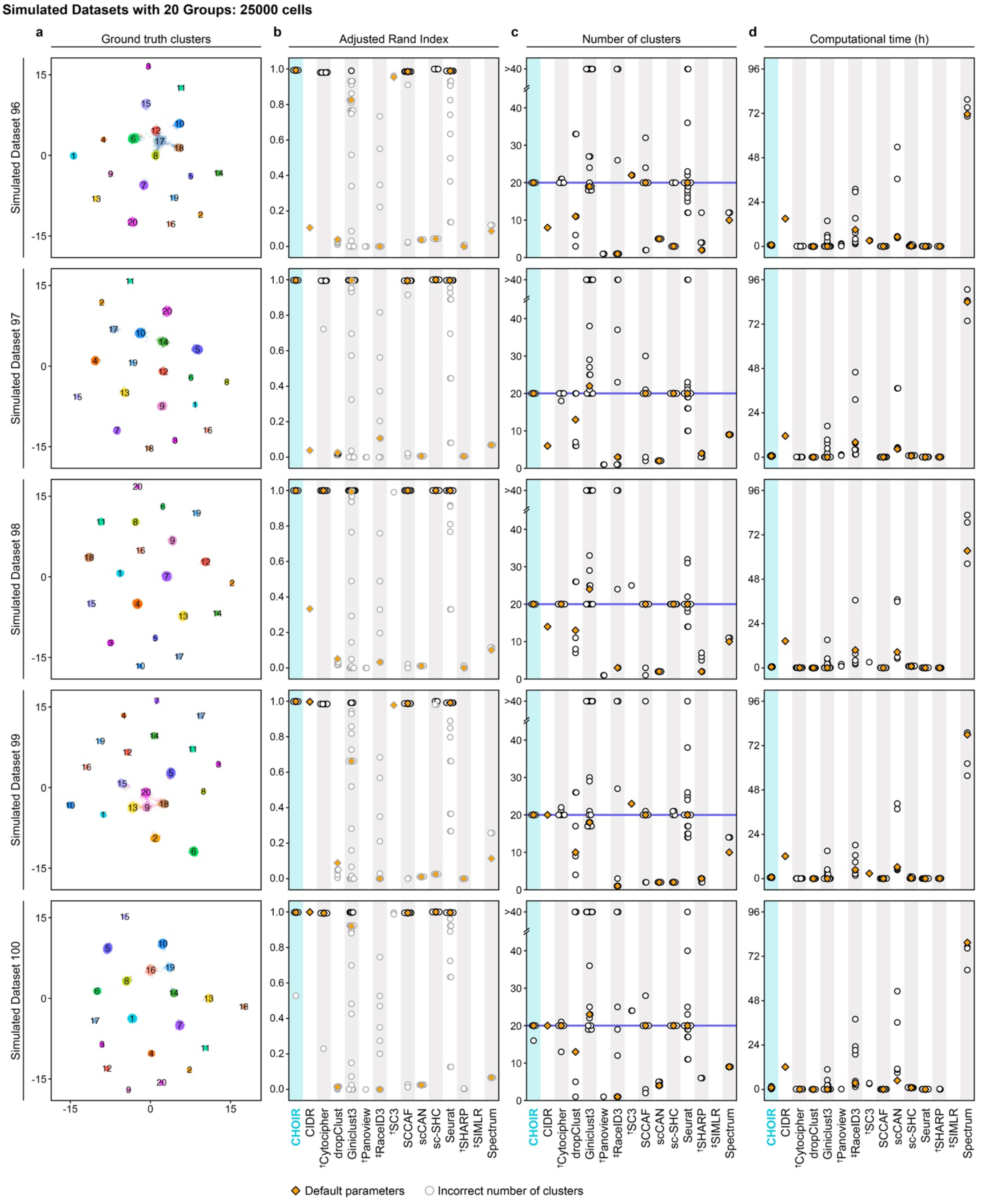
Detailed benchmarking results from simulated datasets with 5–20 ground truth groups. **a**. UMAP embeddings of each dataset colored according to the ground truth groups. **b.** Adjusted Rand Index of each parameter setting tested for each method. Symbols indicate clustering methods for which one or more parameter settings ^†^failed to run or ^‡^did not complete within the maximum allotted runtime of 96 hours for at least one dataset. **c.** Number of clusters of each parameter setting tested for each method. **d.** Computational time of each parameter setting tested for each method.

**Supplementary Fig. 22:**
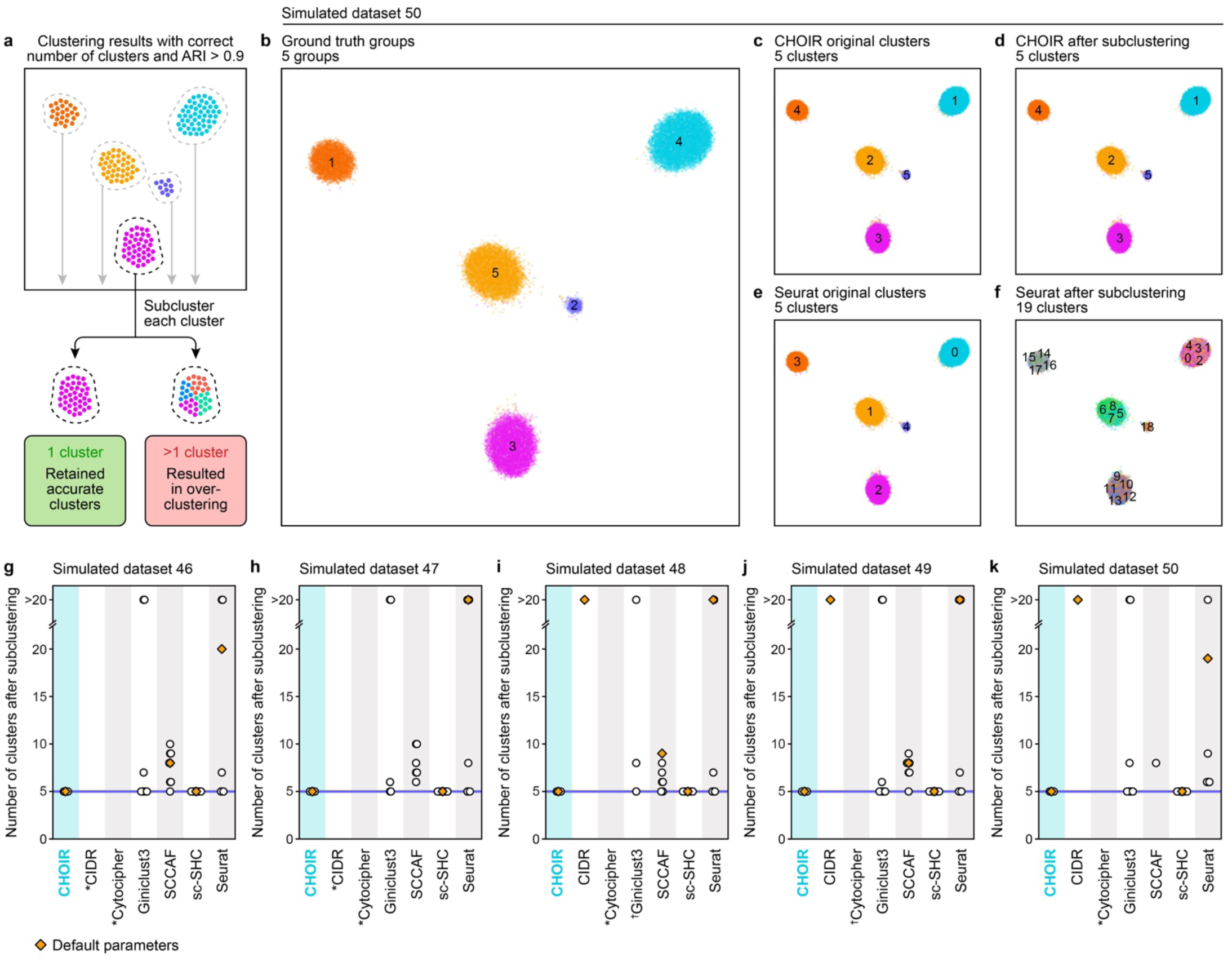
Clusters identified by CHOIR do not require further rounds of subclustering. **a**. Schematic showing the steps of the subclustering analysis. **b–f.** UMAP embedding of Simulated Dataset 50 colored according to the ground truth groups (**b**), the clusters identified using the default parameters of CHOIR (**c**), the clusters identified after subclustering each of the clusters in (**b**) using the default parameters of CHOIR (**d**), the clusters identified using the default parameters of Seurat (**e**), and the clusters identified after subclustering each of the clusters in (**e**) using the default parameters of Seurat (**f**). **g–k.** Dot plots showing the number of clusters obtained after subclustering for each parameter and method combination tested for simulated datasets 46–50. Parameter settings for each cluster method that resulted in the correct number of clusters and achieved an ARI of >0.9 relative to the ground truth groups were tested here. Only seven of the clustering methods tested (CHOIR, CIDR, Cytocipher, GiniClust3, SCCAF, sc-SHC, and Seurat) had any parameter settings meet this condition for any of datasets 46-50. For each such parameter setting, each cluster was subset and subclustered to identify the number of subclusters obtained. Symbols indicate clustering methods for which *no parameter settings identified the correct number of clusters and reached an ARI of >0.9 for the original clustering of the dataset or ^†^one or more parameter settings failed to run.

**Supplementary Fig. 23:**
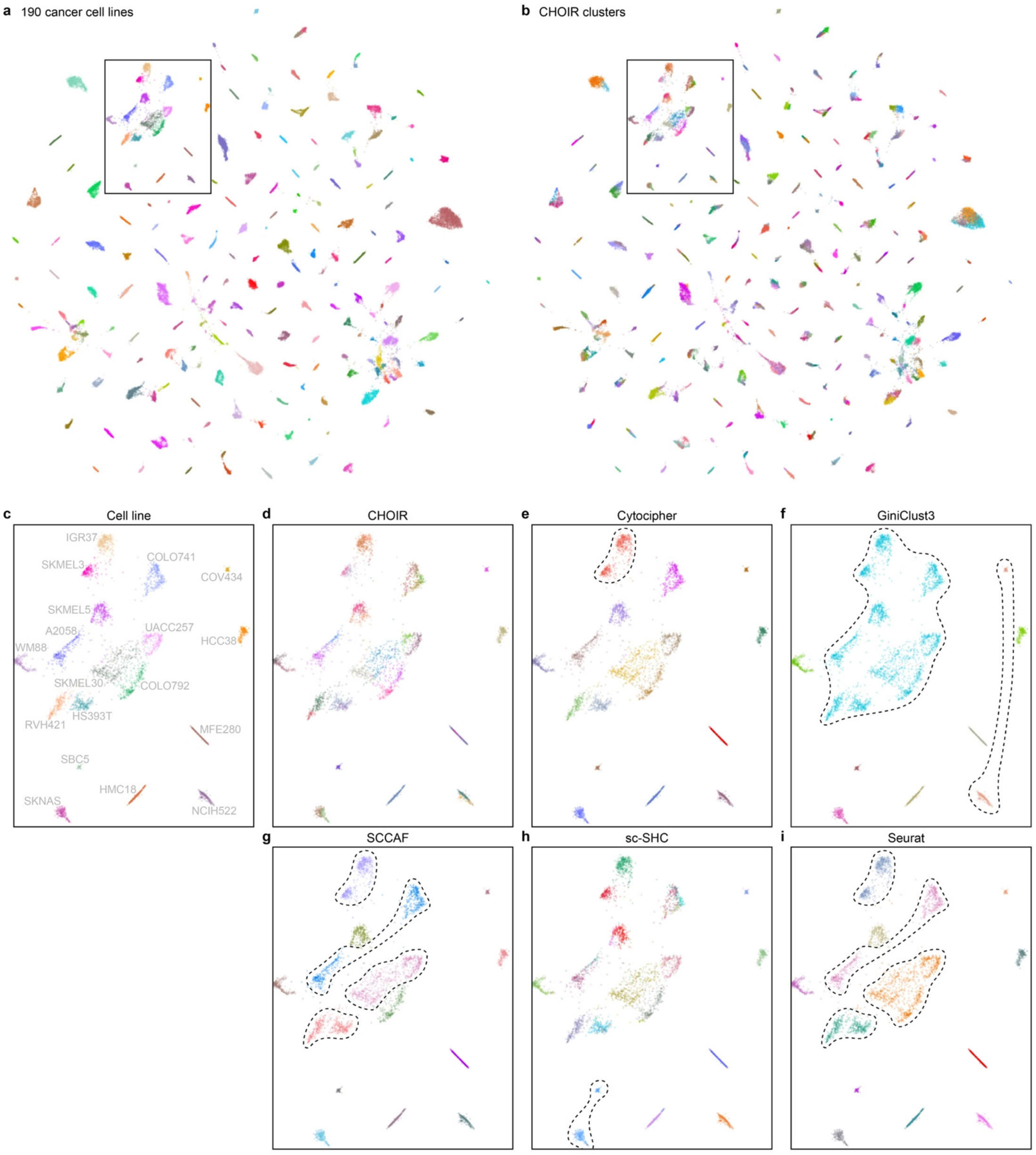
Additional evidence that CHOIR prevents underclustering in an scRNA-seq dataset consisting of pooled cell lines. **a–b**. UMAP embedding of a pooled cancer cell line scRNA-seq dataset from Kinker et al. 2020 consisting of 48,879 cells, colored according to the 190 cancer cell lines (**a**) and the clusters identified using the default parameters of CHOIR (**b**). **b–h.** UMAP embeddings showing the cells within the highlighted rectangle in (**a–b**) colored by cell line (**c**) and the clusters identified by CHOIR (**d**), Cytocipher (**e**), GiniClust3 (**f**), SCCAF (**g**), sc-SHC (**h**), and Seurat (**i**), using the default parameters for each method. Dashed lines indicate instances where the clustering method failed to distinguish individual cell lines, resulting in multiple cell lines grouped within a single cluster.

**Supplementary Fig. 24:**
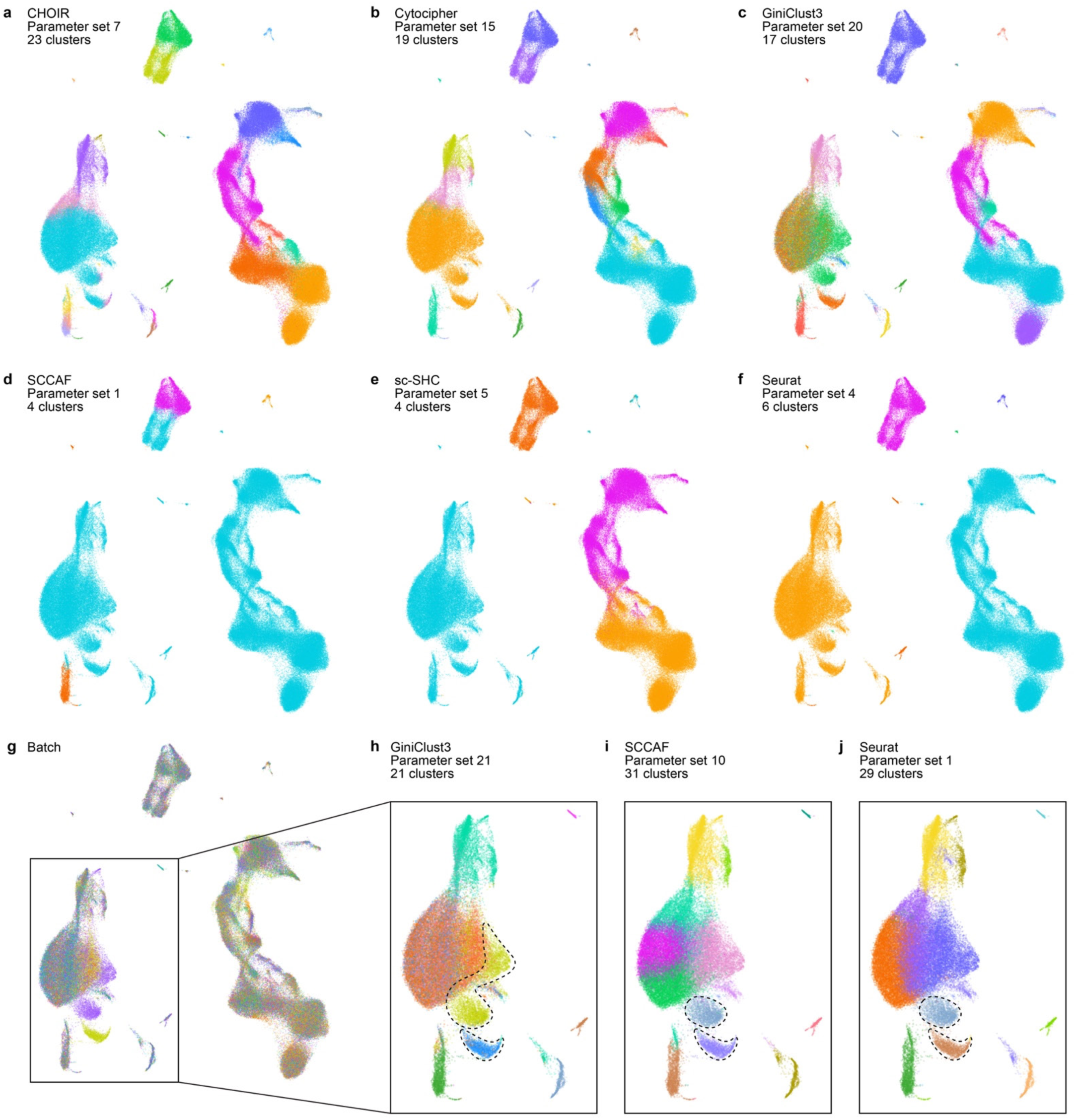
Comparison of results obtained with different clustering methods on Hao et al. 2021 CITE-seq data. **a–f**. UMAP embeddings for the Hao et al. 2021 dataset colored according to the clusters identified by the parameter settings for CHOIR (**a**), Cytocipher (**b**), GiniClust3 (**c**), SCCAF (**d**), sc-SHC (**e**), and Seurat (**f**) that maximized the number of clusters while preventing overclustering as shown in Fig. 4g. **g.** UMAP embedding for the Hao et al. 2021 dataset colored according to batch. The highlighted box identifies a subset of cells where batch effects are observed. **h–j.** UMAP embeddings showing instances in which overclustering was due to confounding by batch for GiniClust3 (**h**), SCCAF (**i**), and Seurat (**j**). Dashed lines indicate clusters confounded by batch.

**Supplementary Fig. 25:**
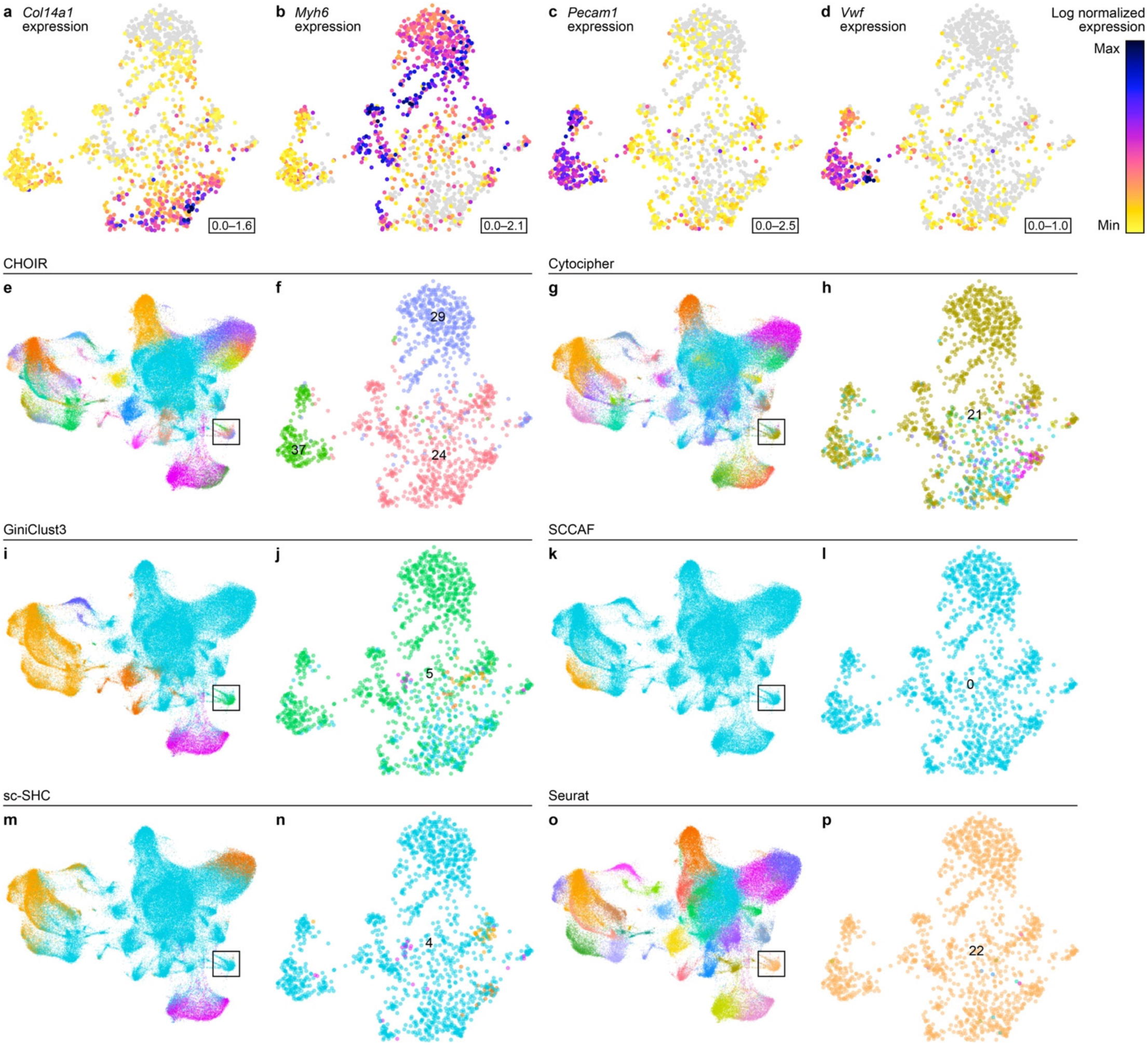
CHOIR identifies three functionally distinct cell types of the embryonic heart. **a–d**. UMAP embedding of the Srivatsan et al. 2021 whole mouse embryo sci-Space dataset computed on the subsetted dimensionality reduction of parent cluster P4 identified using the default parameters in CHOIR, colored according to the expression levels of connective tissue marker gene *Col14a1* (**a**), cardiomyocyte marker gene *Myh6* (**b**), and endothelial cell marker genes *Pecam1* (**c**) and *Vwf* (**d**). **e–p.** UMAP embeddings of all cells (**e, g, i, k, m, o**) and heart cells (**f, h, j, l, n, p**) colored according to the clusters identified by the default parameters of CHOIR (**e–f**), Cytocipher (**g–h**), GiniClust3 (**i–j**), SCCAF (**k–l**), sc-SHC (**m–n**), and Seurat (**o–p**).

**Supplementary Fig. 26:**
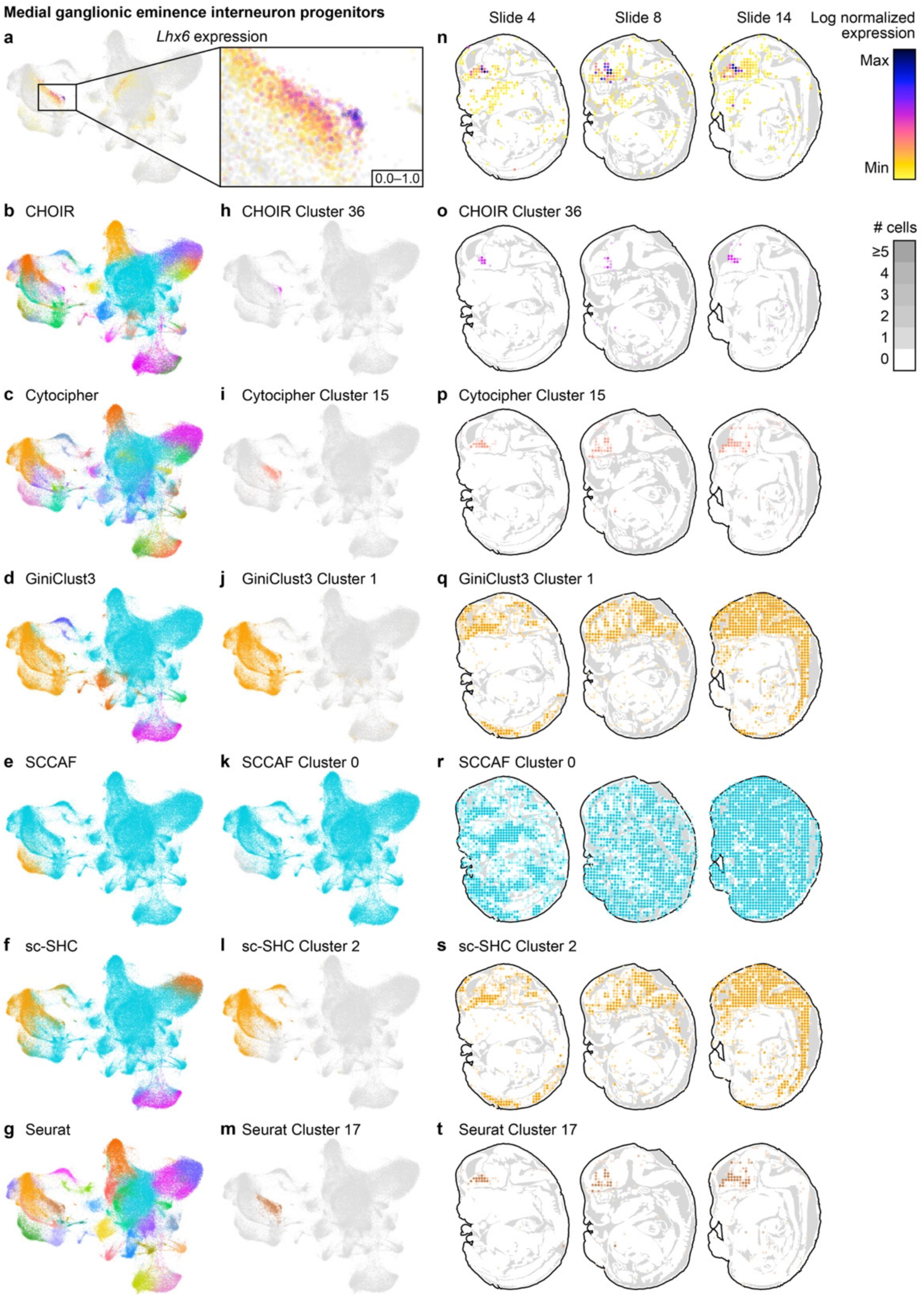
CHOIR cluster 36 represents the pool of medial ganglionic eminence interneuron progenitors characterized by high expression of *Lhx6*. **a–g**. UMAP embedding of the Srivatsan et al. 2021 whole mouse embryo sci-Space dataset, colored according to the expression level of the medial ganglionic eminence interneuron progenitor marker gene *Lhx6* (**a**), or the clusters identified by the default parameters of CHOIR (**b**), Cytocipher (**c**), GiniClust3 (**d**), SCCAF (e), sc-SHC (**f**), or Seurat (**g**). A zoom in of *Lhx6* expression is shown to the right of panel (**a**). **h–m.** UMAP embeddings of the Srivatsan et al. 2021 whole mouse embryo sci-Space dataset, colored by the cluster shown in panels (**a–g**) that harbors the *Lhx6*-expressing medial ganglionic eminence interneuron progenitors for each of the methods indicated. **n**. Distribution of medial ganglionic eminence interneuron progenitor marker gene *Lhx6* in all sections with >25 cells belonging to CHOIR cluster 36 (medial ganglionic eminence interneuron progenitors). **o–t.** Distribution of the method-specific cluster shown to the left in panels (**h–m**) across all sections shown in panel (**n**). The darker the shade of the cluster color, the more cells of the respective cluster were detected at the indicated location.

**Supplementary Fig. 27:**
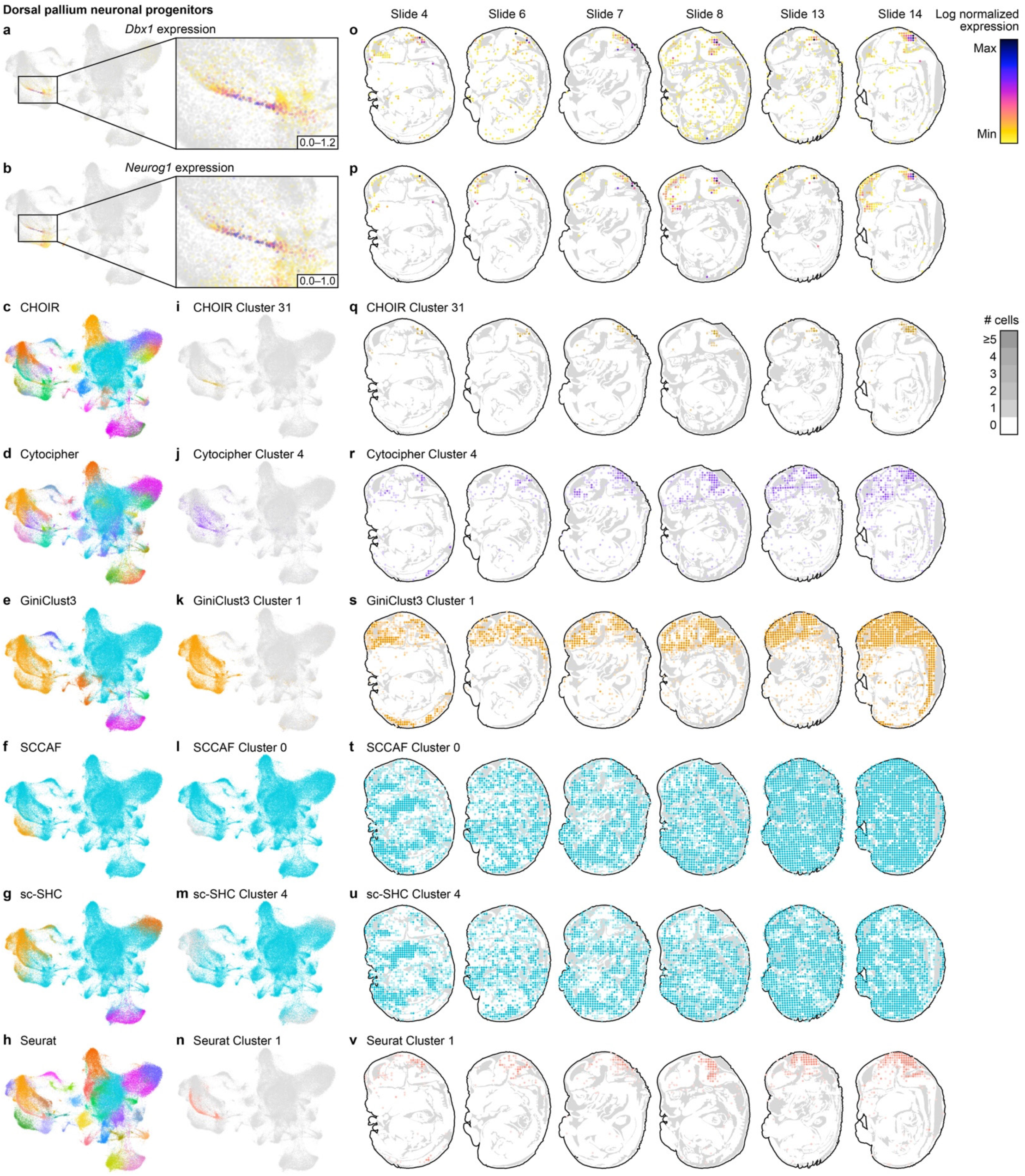
CHOIR cluster 31 represents a neuronal progenitor pool of the spatially restricted E14 dorsal pallium characterized by high expression of *Dbx1* and *Neurog1*. **a–h**. UMAP embeddings of the Srivatsan et al. 2021 whole mouse embryo sci-Space dataset, colored according to the expression levels of dorsal pallium neuronal progenitor marker gene *Dbx1* (**a**) and *Neurog1* (**b**), or the clusters identified by the default parameters of CHOIR (**c**), Cytocipher (**d**), GiniClust3 (**e**), SCCAF (**f**), sc-SHC (**g**), or Seurat (**h**). A zoom in of *Dbx1* and *Neurog1* expression is shown to the right of panels (**a–b**). **i–n.** UMAP embeddings of the Srivatsan et al. 2021 whole mouse embryo sci-Space dataset, colored by the cluster shown in panels (**a–h**) that harbors the *Dbx1– and Neurog1*-expressing dorsal pallium neuronal progenitors for each of the methods indicated. **o–p.** Distribution of dorsal pallium neuronal progenitor marker genes *Dbx1* (**o**) and *Neurog1* (**p**) in all sections with >25 cells belonging to CHOIR cluster 31 (dorsal pallium neuronal progenitors). **q–v.** Distribution of the method-specific cluster shown to the left in panels (**i–n**) across all sections shown in panels (**o–p**). The darker the shade of the cluster color, the more cells of the respective cluster were detected at the indicated location.

**Supplementary Fig. 28:**
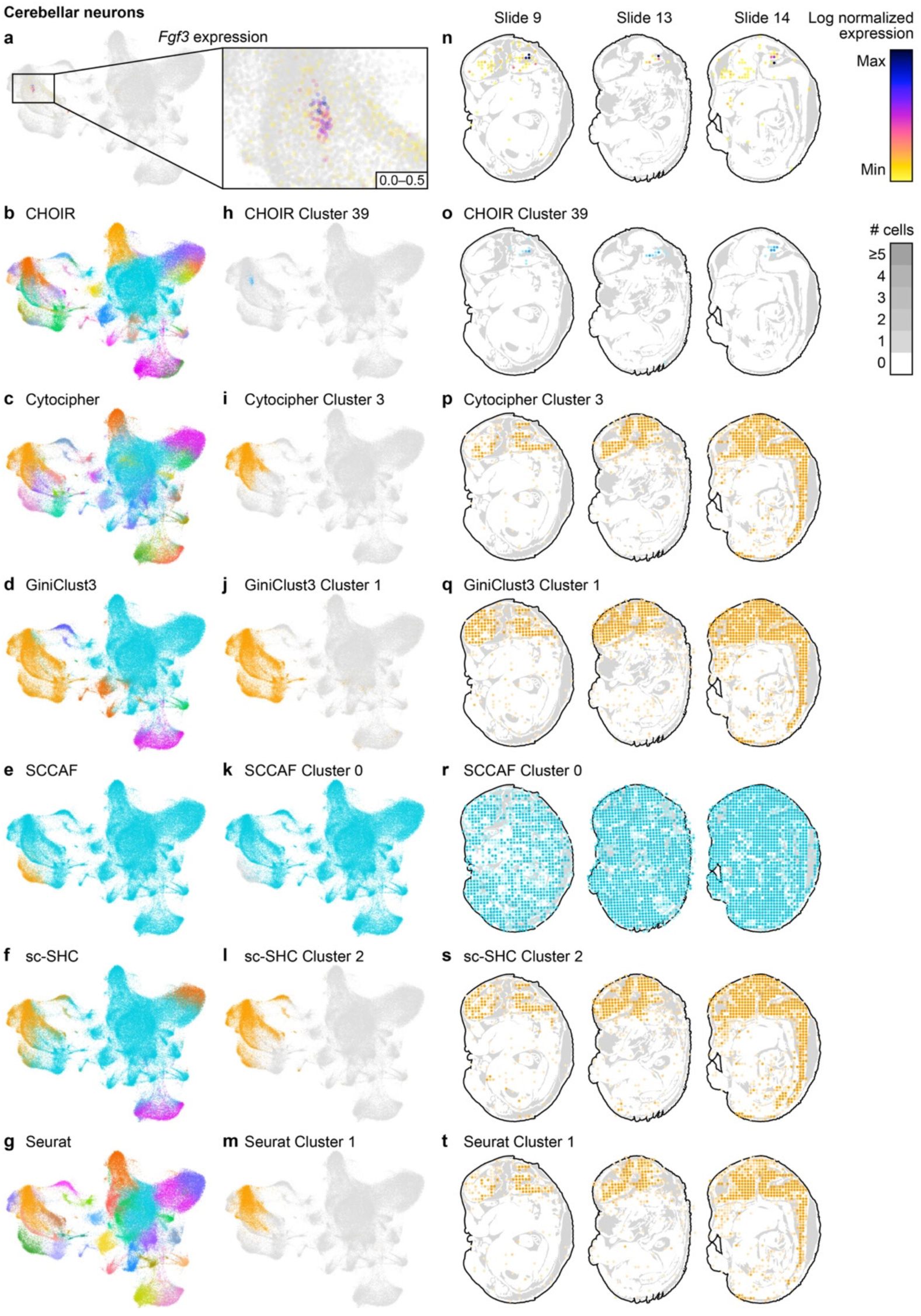
CHOIR cluster 39 represents a population of neurons localized within the developing cerebellum characterized by high expression of *Fgf3*. **a–g**. UMAP embedding of the Srivatsan et al. 2021 whole mouse embryo sci-Space dataset, colored according to the expression level of developing cerebellum marker gene *Fgf3* (**a**), or the clusters identified by the default parameters of CHOIR (**b**), Cytocipher (**c**), GiniClust3 (**d**), SCCAF (**e**), sc-SHC (**f**), or Seurat (**g**). A zoom in of *Fgf3* expression is shown to the right of panel (**a**). **h–m.** UMAP embeddings of the Srivatsan et al. 2021 whole mouse embryo sci-Space dataset, colored by the cluster shown in panels (**a–g**) that harbors the *Fgf3*-expressing developing cerebellum neurons for each of the methods shown to the left. **n.** Distribution of developing cerebellum marker gene *Fgf3* in all sections with >25 cells belonging to CHOIR cluster 39 (developing cerebellum neurons). **o–t.** Distribution of the method-specific cluster shown to the left in panels (**h–m**) across all sections shown in panel (**n**). The darker the shade of the cluster color, the more cells of the respective cluster were detected at the indicated location.

**Supplementary Fig. 29:**
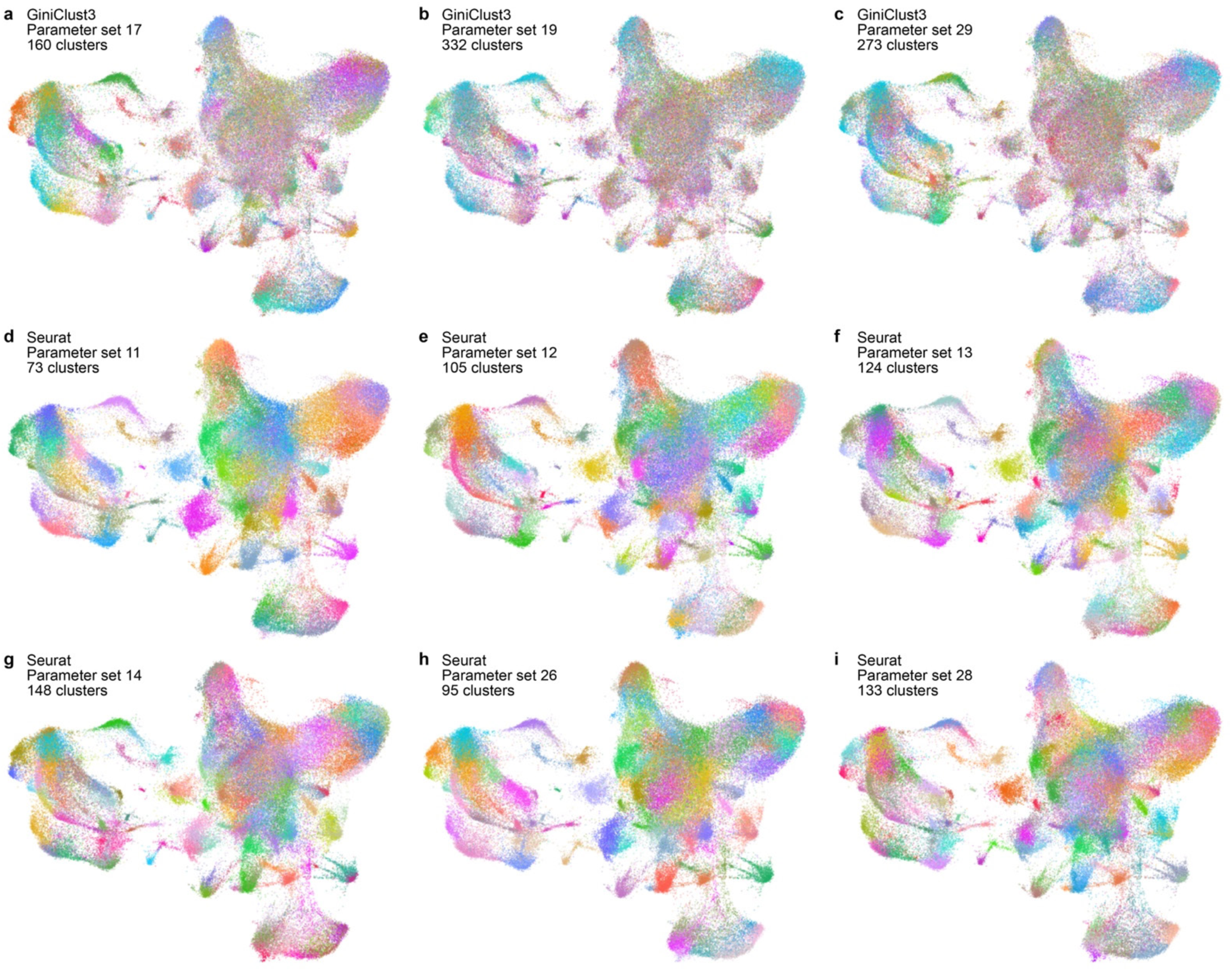
Non-default parameters for GiniClust3 and Seurat that identified certain anatomically restricted cell types resulted in severe overclustering of other cell types. **a–i**. UMAP embeddings of the Srivatsan et al. 2021 whole mouse embryo sci-Space dataset colored according to the clusters identified by GiniClust3 (**a–c**) or Seurat (**d–i**) using non-default parameters. Although these non-default parameters identified at least 3 of the 7 functionally distinct cell types in the heart or specific brain regions that were identified by CHOIR’s default parameters, they resulted in severe overclustering of other cell types.

